# KLF10 integrates circadian timing and sugar signaling to coordinate hepatic metabolism

**DOI:** 10.1101/2020.12.22.423999

**Authors:** Anthony A. Ruberto, Aline Gréchez-Cassiau, Sophie Guérin, Luc Martin, Johana S. Revel, Mohamed Mehiri, Malayannan Subramaniam, Franck Delaunay, Michèle Teboul

**Affiliations:** Université Côte d’Azur, CNRS, Inserm, iBV, 06108 Nice, France; Université Côte d’Azur, CNRS, Institut de Chimie de Nice, UMR 7272, 06108 Nice, France; Department of Biochemistry and Molecular Biology, Mayo Clinic, Rochester, MN 55905, USA; Department of Parasites and Insect Vectors, Institut Pasteur, 75015 Paris, France

## Abstract

The mammalian circadian timing system and metabolism are highly interconnected, and disruption of this coupling is associated with negative health outcomes. Krüppel-like factors (KLFs) are transcription factors that govern metabolic homeostasis in various organs. Many KLFs show a circadian expression in the liver. Here, we show that the loss of the clock-controlled KLF10 in hepatocytes results in extensive reprogramming of the mouse liver circadian transcriptome, which in turn, alters the temporal coordination of pathways associated with energy metabolism. We also show that glucose and fructose induce *Klf10,* which helps mitigate glucose intolerance and hepatic steatosis in mice challenged with a sugar beverage. Functional genomics further reveal that KLF10 target genes are primarily involved in central carbon metabolism. Together, these findings show that in the liver, KLF10 integrates circadian timing and sugar metabolism related signaling, and serves as a transcriptional brake that protects against the deleterious effects of increased sugar consumption.

## Introduction

The mammalian circadian timing system aligns most biological processes with the Earth’s light/dark (LD) cycle to ensure optimal coordination of physiology and behavior over the course of the day (Bass and Lazar, 2016). At the organism level, circadian clocks are organized hierarchically: at the top a central pacemaker located in the suprachiasmatic nuclei of the hypothalamus that receives the light input and in turn coordinates peripheral clocks *via* internal synchronizers including glucocorticoids and body temperature (Saini et al., 2011). The molecular mechanism underlying circadian clocks is an oscillatory gene network present in virtually all cells which translates the external time information into optimally phased rhythms in chromatin remodeling, gene expression, posttranslational modification and metabolites production (Dyar et al., 2018; Mauvoisin and Gachon, 2019; Rijo-Ferreira and Takahashi, 2014; Robles et al., 2017, 2014; Weidemann et al., 2018; Zhang et al., 2014).

The extensive circadian regulation of processes linked to metabolic homeostasis, as well as the mechanisms by which cellular metabolism feeds back to core clock components, have been highlighted by functional genomics studies (Panda, 2016; Peek et al., 2013; Reinke and Asher, 2019; Sinturel et al., 2020). In mice, the disruption of clock genes leads to multiple metabolic abnormalities, including impaired insulin release, glucose intolerance, insulin resistance, hepatic steatosis and obesity (Bugge et al., 2012; Jacobi et al., 2015; Perelis et al., 2015); conversely, diet-induced metabolic imbalance impairs circadian coordination in part by reprogramming the circadian transcriptome (Eckel-Mahan et al., 2013; Peek et al., 2013). Experimental and epidemiological data support a similar reciprocal relationship between circadian misalignment or disruption and metabolic disorders in humans (Qian and Scheer, 2016; Stenvers et al., 2019).

The liver is a major metabolic organ in which circadian gene expression is regulated directly by core clock and clock-controlled transcription factors among which nuclear hormone receptors including REV-ERBs, PPARs, GR, FXR, SHP, play a prominent role (Mukherji et al., 2019). Krüppel-like factors (KLFs) form another family of transcription regulators, a majority of which has been involved the physiology of metabolic organs including the liver (Hsieh et al., 2019). Many of these KLFs are also clock-controlled genes in mouse liver (Ceglia et al., 2018; Yoshitane et al., 2014). This suggests KLFs as additional factors for the circadian regulation of hepatic metabolism. KLF15 was for instance shown to control the circadian regulation of nitrogen metabolism and bile acid production (Han et al., 2015; Jeyaraj et al., 2012). KLF10, first described as a TGF-β induced early gene in human osteoblasts and pro-apoptotic factor in pancreatic cancer (Subramaniam et al., 1995; Tachibana et al., 1997), has been further identified as a clock-controlled and glucose-induced transcription factor that helps suppress hepatic gluconeogenesis (Guillaumond et al., 2010; Hirota et al., 2002). In *Drosophila*, the KLF10 ortholog *Cabut*, has a similar role in sugar metabolism and circadian regulation (Bartok et al., 2015), which suggests the function of the transcription factor is evolutionary conserved.

A better understanding of the role of KLF10 in hepatic metabolism may have important translational implications in the context of the rising prevalence of nonalcoholic fatty liver disease (NAFLD) and fructose overconsumption (Jensen et al., 2018). In this study, using a hepatocyte-specific *Klf10* knockout mouse model, we show that KLF10 is required for the temporal coordination of various biological pathways associated with energy metabolism, and that it has a protective role in shielding mice from adverse effects associated with sugar overload.

## Results

### Hepatocyte-specific deletion of KLF10 reprograms the liver circadian transcriptome

To understand the role of the circadian transcription factor KLF10 in hepatic metabolism, we generated a conditional *Klf10* knockout mouse model by crossing *Klf10*^*flox/flox*^ mice (Weng et al., 2017) with a mouse line expressing an inducible CreER^T2^ recombinase under the control of the endogenous serum albumin (SA) promoter (Schuler et al., 2004) (Figure 1A). Upon tamoxifen treatment of *Klf10*^*flox/flox*^, *SA*^*+/CreERT2*^ mice, we obtained *a* deletion of *Klf10* specifically in hepatocytes (*Klf10*^*Δhep*^). Both qPCR and immunoblot analyses confirmed the targeted reduction of the transcription factor in the liver of *Klf10*^*Δhep*^ mice, with residual expression in this organ coming from non-parenchymal cells (Figures 1B and 1C). As expected, we detected a robust circadian oscillation of *Klf10* mRNA in the liver of *Klf10*^*flox/flox*^ mice, with peak expression of the transcript occurring just before the LD transition while in *Klf10^Δhep^* mice, rhythmic expression of *Klf10* was absent (Figure 1D). Body weight, food intake and food seeking behavior did not differ between genotypes, indicating that post-natal hepatocyte-specific deletion of KLF10 did not grossly affect the physiology and behavior of unchallenged adult *Klf10*^*Δhep*^ mice (Figures S1A and S1B).

**Figure 1.**
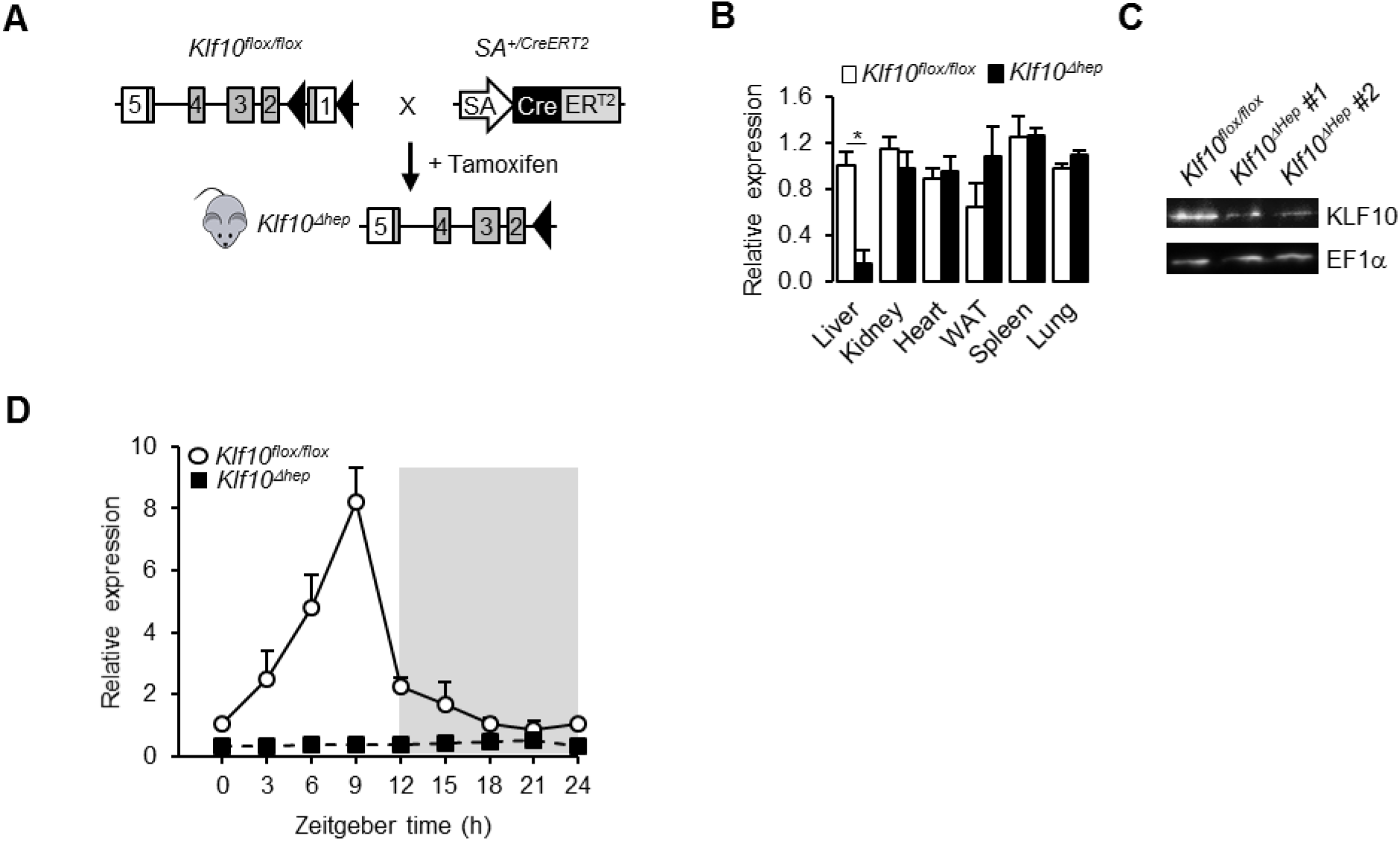
Genetic disruption of *Klf10* in mouse hepatocytes. (A) Schematic illustrating the strategy for generating of *Klf10*^*Δhep*^ mice. (B) Relative gene expression of *Klf10* across various tissue in *Klf10*^*flox/flox*^ and *Klf10*^*Δhep*^ mice as measured at ZT9 by RT-qPCR (mean ± SEM, n = 3-5). (C) Immunoblot showing KLF10 protein abundance in liver extracts from *Klf10*^*flox/flox*^ and *Klf10*^*Δhep*^ mice at ZT9. EF1α was used as loading control. (D) 24h gene expression profiles of hepatic *Klf10* mRNA measured every 3 hours in *Klf10*^*flox/flox*^ and *Klf10*^*Δhep*^ mice (mean ± SEM, n = 3 mice per time point). Statistics: Non parametric Wilcoxon test. * p < 0.05. See also Figure S1.

We next performed RNA sequencing (RNA-seq) on livers collected every three hours across the 24h day from *Klf10*^*flox/flox*^ and *Klf10*^*Δhep*^ mice entrained to a 12h:12h LD cycle (Figure 2A). Using MetaCycle (Wu et al., 2016), we found that ~12% of genes (1,664) displayed circadian expression in *Klf10*^*flox/flox*^ mice (Figure 2B; Table S1), a percentage which is consistent with previous reports (Greenwell *et al.*, 2019; Zhang *et al.*, 2014). In *Klf10*^*Δhep*^mice, we detected a similar percentage of genes with circadian expression (~12%; 1,633) (Figure 2B; Table S1). The phase distributions and relative amplitudes of rhythmically expressed genes in *Klf10*^*flox/flox*^ and *Klf10*^*Δhep*^ mice were similar (Figures 2C and 2D). Furthermore, the 24h expression profiles of the core clock genes in *Klf10*^*Δhep*^ mice were unchanged (Figure S2A), suggesting that hepatocyte KLF10 is not required for normal core clock function in the liver. Despite these similarities, less than 40% of genes (602) displayed rhythmic expression in both genotypes (Figures 2E and 2F). Interestingly, we observed rhythmicity of 1,031 transcripts exclusively in *Klf10*^*Δhep*^ mice, indicating that the deletion of KLF10, a known transcriptional repressor, results in *de novo* oscillation of otherwise non-rhythmically expressed transcripts (Figures 2E and 2F).

**Figure 2.**
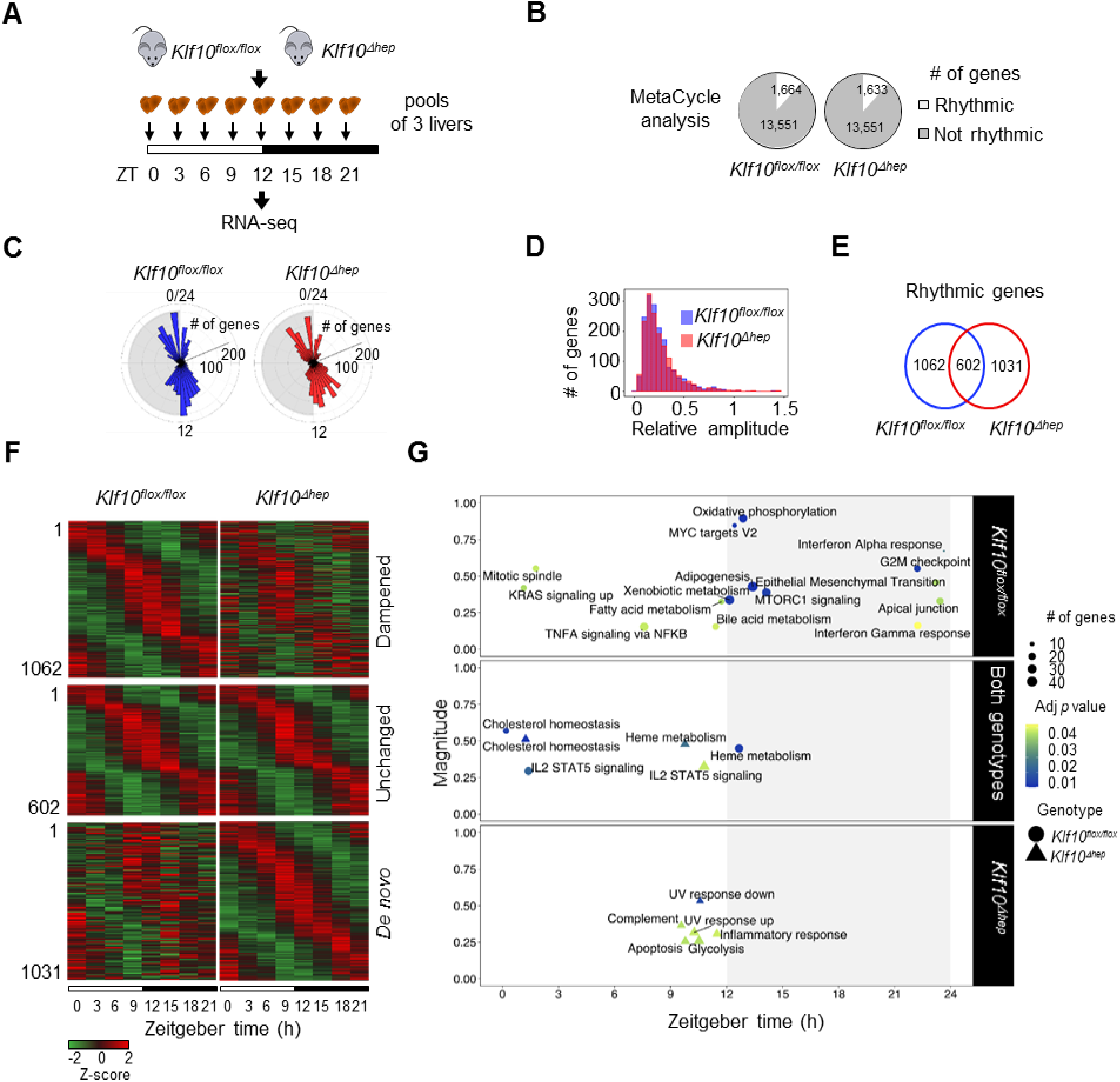
Deletion of hepatocyte KLF10 alters the circadian transcriptome in the liver. (A) Schematic illustrating the workflow used to assess the circadian transcriptomes of livers in *Klf10*^*flox/flox*^ and *Klf10*^*Δhep*^ mice. (B) Number of rhythmic transcripts detected in the livers from *Klf10*^*flox/flox*^ and *Klf10*^*Δhep*^ mice. (C) Phase distributions of rhythmic transcripts in the livers of *Klf10*^*flox/flox*^ and *Klf10*^*Δhep*^ mice. (D) Relative amplitudes of rhythmic transcripts in the livers of *Klf10*^*flox/flox*^ and *Klf10*^*Δhep*^ mice. (E) Number of unique and overlapping rhythmic transcripts in *Klf10*^*flox/flox*^ and *Klf10*^*Δhep*^ mice. (F) Heatmap showing the expression profiles of oscillating transcripts in the livers of *Klf10*^*flox/flox*^ and *Klf10*^*Δhep*^ mice (pools of 3 livers per time point). (G) Magnitudes and phases of enriched biological processes identified by PSEA in *Klf10*^*flox/flox*^ mice only (top), in *Klf10*^*Δhep*^ mice only (bottom), and in both genotypes (middle). Statistics: MetaCycle, significance threshold, *p* < 0.05; PSEA; Kuiper test, significance threshold, q < 0.01. See also Figure S2 and Tables S1 and S2.

To better understand the changes in the circadian transcriptome associated with the deletion of *Klf10* in hepatocytes, we assessed the temporal coordination of biologically related transcripts in *Klf10*^*flox/flox*^ and *Klf10*^*Δhep*^ mice. After performing Phase Set Enrichment Analysis (PSEA) (Zhang et al., 2016), we found a decrease in the total number of temporally enriched gene sets in *Klf10*^*Δhep*^ mice (Figure 2G). Specifically, gene sets associated with processes such as oxidative phosphorylation, fatty acid metabolism, and mTORC1 signaling peaking around the LD transition in *Klf10*^*flox/flox*^ mice were absent in *Klf10*^*Δhep*^ mice (Figure 2G and Table S2). Despite the global reduction in temporally coordinated transcripts, *Klf10*^*Δhep*^ mice acquired *de novo* temporal enrichment of transcripts associated with glycolysis and cellular stress processes such as apoptosis, UV response and inflammation (Figure 2G, Table S2). The alterations of biological pathways at the transcript level in *Klf10*^*Δhep*^ mice prompted us to assess whether any changes associated with these pathways would be visible at the physiological level. Given the scope of our research question, we focused primarily on processes linked to metabolism. We found that *Klf10*^*Δhep*^ hepatocytes displayed increased glucose uptake capacity (Figure S2B), and that *Klf10*^*Δhep*^ mice had decreased glycogen levels during their resting phase (Figure S2C), suggesting that the loss of *Klf10* in hepatocytes leads to subtle physiological changes in carbohydrate processing at both the cellular and tissue level. Together, these findings indicate that dysregulation of the hepatic circadian transcriptome in *Klf10*^*Δhep*^ mice results in the rewiring of various pathways related to energy metabolism, which in turn may compromise the ability of these animals to process energy substrates.

### Hepatocyte KLF10 minimizes the adverse metabolic effects associated with a high sugar diet

Consumption of glucose and fructose in the form of sugar-sweetened beverages is linked to negative health outcomes (Softic et al., 2020). While previous studies have shown that glucose is a potent inducer of *Klf10* expression in primary hepatocytes (Guillaumond et al., 2010; Iizuka et al., 2011), the effect of fructose is unknown. We now show that hepatocytes treated with 5 mM fructose induces *Klf10* to the same extent as high (25 mM) glucose; while the combination of high glucose and fructose has an additive effect (Figure 3A). Given these results in hepatocytes, combined with transcriptional changes occurring in *Klf10*^*Δhep*^ mice, we sought to better understand the role of KLF10 in mediating the effect of the two dietary sugars at the organism level. *Klf10*^*flox/flox*^ and *Klf10*^*Δhep*^ mice were given *ad libitum* access to a chow diet with water (chow) or a chow diet with sugar-sweetened water (chow + SSW) for 8 weeks (Figure 3B). Similar to the response we observed in hepatocytes, the chow + SSW diet induced hepatic *Klf10* expression, when measured during the animals’ feeding phase (ZT15) (Figure 3C). Mice on the chow + SSW diet adapted their feeding behavior, with less calories coming from the chow pellets. As a result, their total calorie intake was unchanged relative to mice fed the chow diet (Figures S3A). After eight weeks on the chow + SSW diet, mice from both genotypes displayed a significant increase in body mass and epididymal fat pad mass (Figures 3D and 3E), while only *Klf10*^*Δhep*^ mice displayed an increase in liver mass (Figure 3F). Chow + SSW diet increased liver triglycerides content, and histologic signs of steatosis were evident in both genotypes; however, to a greater extent in *Klf10*^*Δhep*^ mice (Figure 3G). Despite the greater increase in liver triglycerides in *Klf10*^*Δhep*^ mice, no difference in circulating triglycerides was detected between genotypes (Figure S3B).

**Figure 3.**
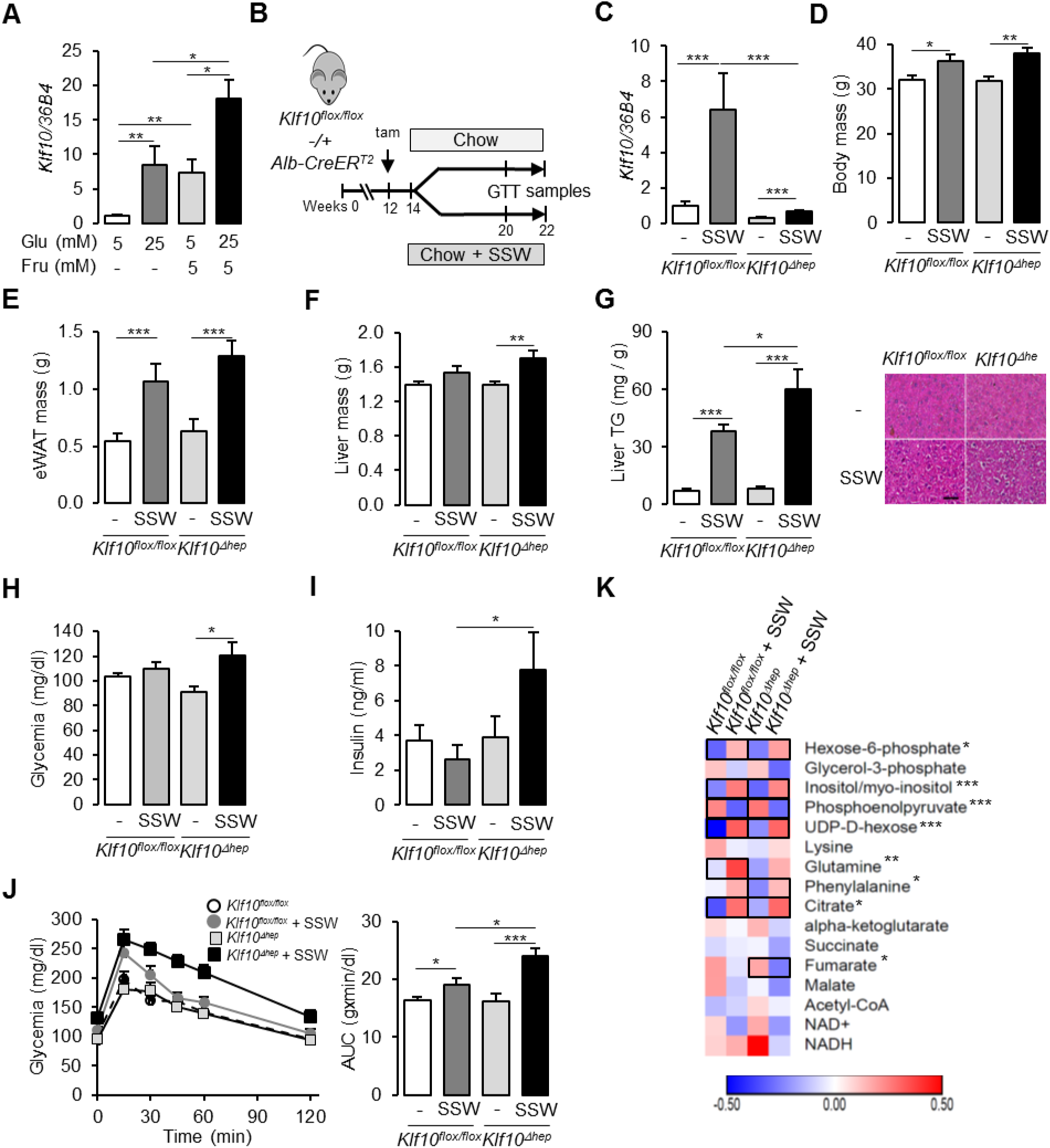
Loss of hepatocyte KLF10 exacerbates the adverse effects associated with increased sugar consumption. (A) Response of *Klf10* in primary mouse hepatocytes challenged with glucose and/or fructose *in vitro* (mean ±SEM, n = 6). (B) Schematic illustrating the duration and sampling time points of mice undergoing the SSW paradigm. (C) Gene expression of *Klf10* in livers of *Klf10*^*flox/flox*^ and *Klf10*^*Δhep*^ mice given a chow or chow + SSW diet (mean ± SEM, n = 4). (D) Body mass of mice given a chow or chow + SSW diet (mean ± SEM, n = 9-12). (E) Epididymal white adipose tissue mass of mice given a chow or chow + SSW diet (mean ± SEM, n = 9-12). (F) Liver mass of mice given a chow or chow + SSW diet (mean ± SEM, n = 8-12). (G) Liver triglyceride content (mean ± SEM, n = 6-10) (left) and representative images of liver histology (right) in *Klf10*^*flox/flox*^ and *Klf10*^*Δhep*^ mice given a chow or chow + SSW diet (scale bar = 50 μm). (H) Blood glucose in *Klf10*^*flox/flox*^ and *Klf10*^*Δhep*^ mice given a chow or chow + SSW diet (n = 9-12). (I) Insulin levels in *Klf10*^*flox/flox*^ and *Klf10*^*Δhep*^ mice given a chow or chow + SSW diet (mean ± SEM, n = 9-12). (J) Blood glucose levels assessed at regular intervals over a 2 hour period in mice undergoing a glucose tolerance test (GTT) performed at ZT 12 (left) and area under the curve of blood glucose levels over the measurement period (right) (mean ± SEM, n = 9-12). (K) Heatmap showing the normalized concentration of metabolites in the liver of *Klf10*^*flox/flox*^ and *Klf10*^*Δhep*^ mice given a chow or chow + SSW diet (mean, n = 9-12). Significant pairwise comparisons are boxed. Unless otherwise indicated all measurements were performed during the animals’ feeding period, at ZT15. Statistics: Non parametric Kruskal and Wallis test. \**p* < 0.05, \*\**p* < 0.01, *** *p*<0.005. See also Figure S3.

To further profile these mice, we assessed various parameters associated with sugar metabolism during the animals’ early active (feeding) phase (ZT12-15)—a time of day when insulin sensitivity is at its maximum (la Fleur et al., 2001), and when pathways associated with nutrient intake and processing are activated (Greenwell et al., 2019; Vollmers et al., 2009). Compared to *Klf10*^*flox/flox*^ mice, *Klf10*^*Δhep*^ mice on the chow + SSW diet had a significant increase in glycemia, and displayed hyperinsulinemia (Figure 3H and 3I). Furthermore, *Klf10*^*Δhep*^ mice on the chow + SSW diet exhibited greater glucose intolerance compared to *Klf10*^*flox/flox*^ mice on the same diet (Figure 3J). Next we profiled carbohydrate, amino acid and TCA related metabolite levels in the four experimental groups of mice. We found that hexose-6-phosphate, myo-inositol, UDP-D-hexose and citric acid were increased and that phospho-enol-pyruvate was decreased to a similar extent in both *Klf10*^*flox/flox*^ and *Klf10*^*Δhep*^ mice on the chow + SSW diet (Figure 3K). In contrast, glutamine was increased only in *Klf10*^*flox/flox*^ on the chow +SSW diet, while phenylalanine and fumarate were increased and decreased respectively only in *Klf10*^*Δhep*^ mice on the chow + SSW (Figure 3K). These observations reveal subtle changes in amino acid catabolism in mice lacking hepatocyte KLF10.

We next assessed the effect of the chow + SSW diet at the transcriptional level in *Klf10*^*flox/flox*^ and *Klf10*^*Δhep*^ mice by comparing the expression of various gene associated with metabolism. We detected an upregulation of *Slc2a4,* a glucose transporter with known expression in adipose and skeletal muscle (Klip et al., 2019), and normally undetected in the liver (Figure 4A, left). Furthermore, the absence of KLF10 in hepatocytes dramatically increased this upregulation of *Slc2a4* in *Klf10*^*Δhep*^ mice on chow + SSW compared to *Klf10*^*flox/flox*^ mice on the same diet (Figure 4A). In contrast, the chow + SSW diet resulted in similar upregulation of glucose (*Slc2a2)* and fructose (*Slc2a5)* transporters in both *Klf10*^*flox/flox*^ and *Klf10*^*Δhep*^ mice (Figure S4). Upon assessing genes involved in the glucose and fructose metabolism pathway, we found that *Klf10*^*Δhep*^ mice on the chow + SSW diet displayed higher expression of the gene encoding for the rate-limiting glycolytic enzyme encoding *Pklr* compared to *Klf10*^*flox/flox*^ mice on the same diet (Figure 4A). Furthermore, the chow + SSW diet altered expression of genes encoding for enzymes associated with gluconeogenesis. However, we detected an upregulation of *G6Pase* in mice from both genotypes on the chow + SSW diet, whereas the expression of *Pck1*, a direct KLF10 target gene (Guillaumond et al., 2010), was upregulated exclusively in *Klf10*^*Δhep*^ mice (Figure 4B). Hepatocyte-specific deletion of KLF10 also altered the transcriptional response of genes involved in *de novo* lipogenesis, as *Klf10*^*Δhep*^ mice fed the chow + SSW diet displayed a significantly greater induction of the genes encoding for *Acly, Acc1, Fasn, Elovl6, ME, and Thrsp* relative to their *Klf10*^*flox/flox*^ counterparts (Figure 4C).

**Figure 4.**
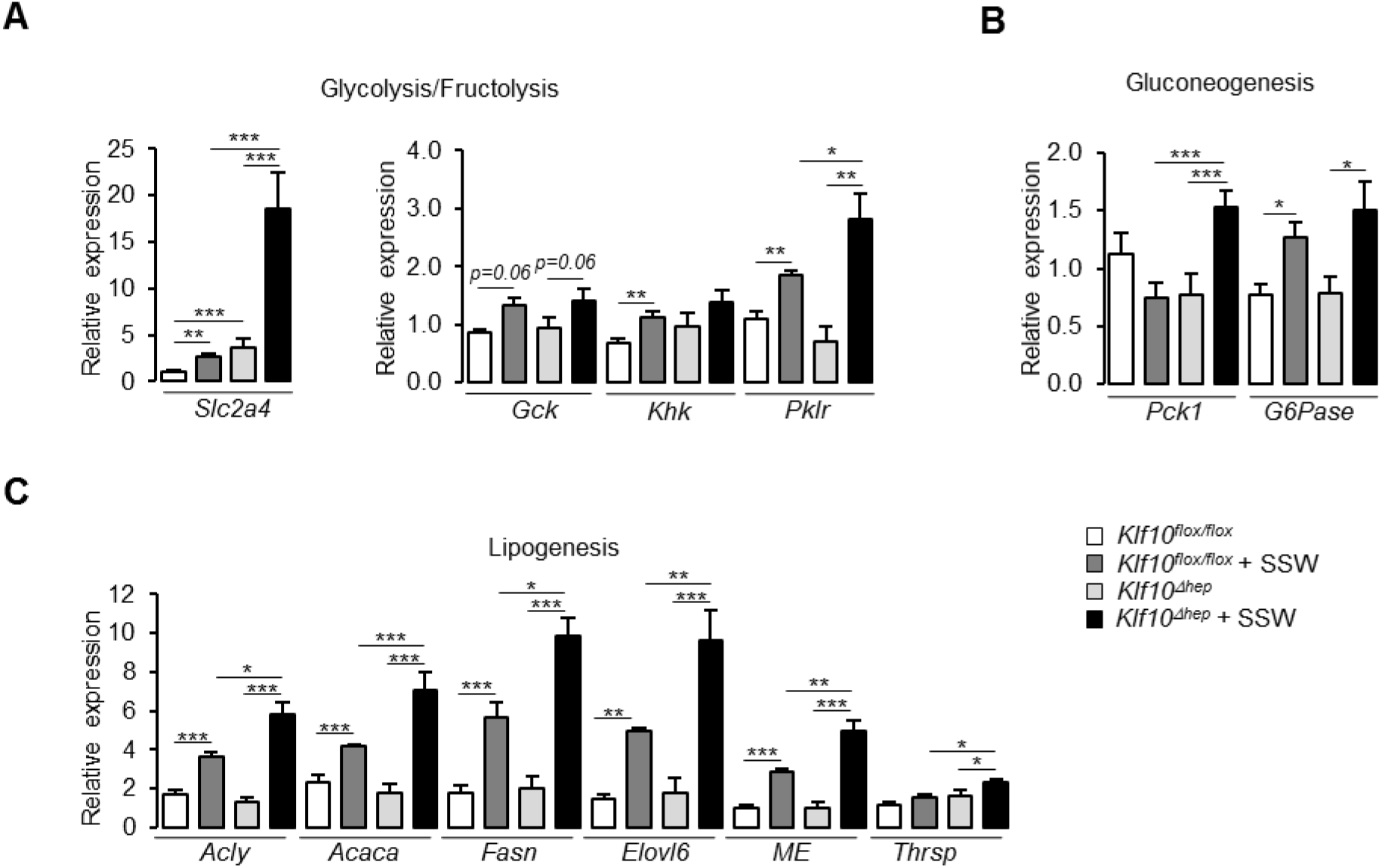
Altered metabolic gene expression in *Klf10*^*Δhep*^ mice challenged with high sugar. (A) Expression profiles of genes associated with glycolysis/fructolysis in the liver of *Klf10*^*flox/flox*^ and *Klf10*^*Δhep*^ mice given a chow or chow + SSW diet (mean ± SEM, n = 6). (B) Expression profiles of genes associated with gluconeogenesis in the liver of *Klf10*^*flox/flox*^ and *Klf10*^*Δhep*^ mice given a chow or chow + SSW diet (mean ± SEM, n = 5-6). (C) Expression profiles of genes associated with lipogenesis in the liver of *Klf10*^*flox/flox*^ and *Klf10*^*Δhep*^ mice given a chow or chow + SSW diet (mean ± SEM, n = 4-6). Statistics: Non parametric Kruskal and Wallis test. \**p* < 0.05, \*\**p* < 0.01, ***, *p*<0.005. See also Figure S4.

Taken together, our physiological, metabolic and molecular data suggests that the induction of KLF10 in response to a dietary sugar challenge serves as a protective mechanism that aids in safeguarding the organism from the detrimental effects associated with excess sugar intake.

### KLF10 is a ‘transcriptional brake’ that fine-tunes sugar signaling in hepatocytes

As the induction of hepatic KLF10 in chow +SSW fed mice helps to minimize the adverse metabolic effects associated with excess dietary sugar intake, we hypothesized that this transcription factor may be involved in the metabolic adaptation of hepatocytes to acute changes in carbohydrate availability. To test this hypothesis in a cell autonomous manner, we treated *Klf10*^*flox/flox*^ and *Klf10*^*Δhep*^ primary hepatocytes with either 5 mM glucose (low glucose; LG) or 25 mM glucose and 5 mM fructose (high glucose and fructose; HGF) for 12 hours (Figure 5A and 5B) and performed RNA-seq to profile the transcriptomes under each condition.

**Figure 5.**
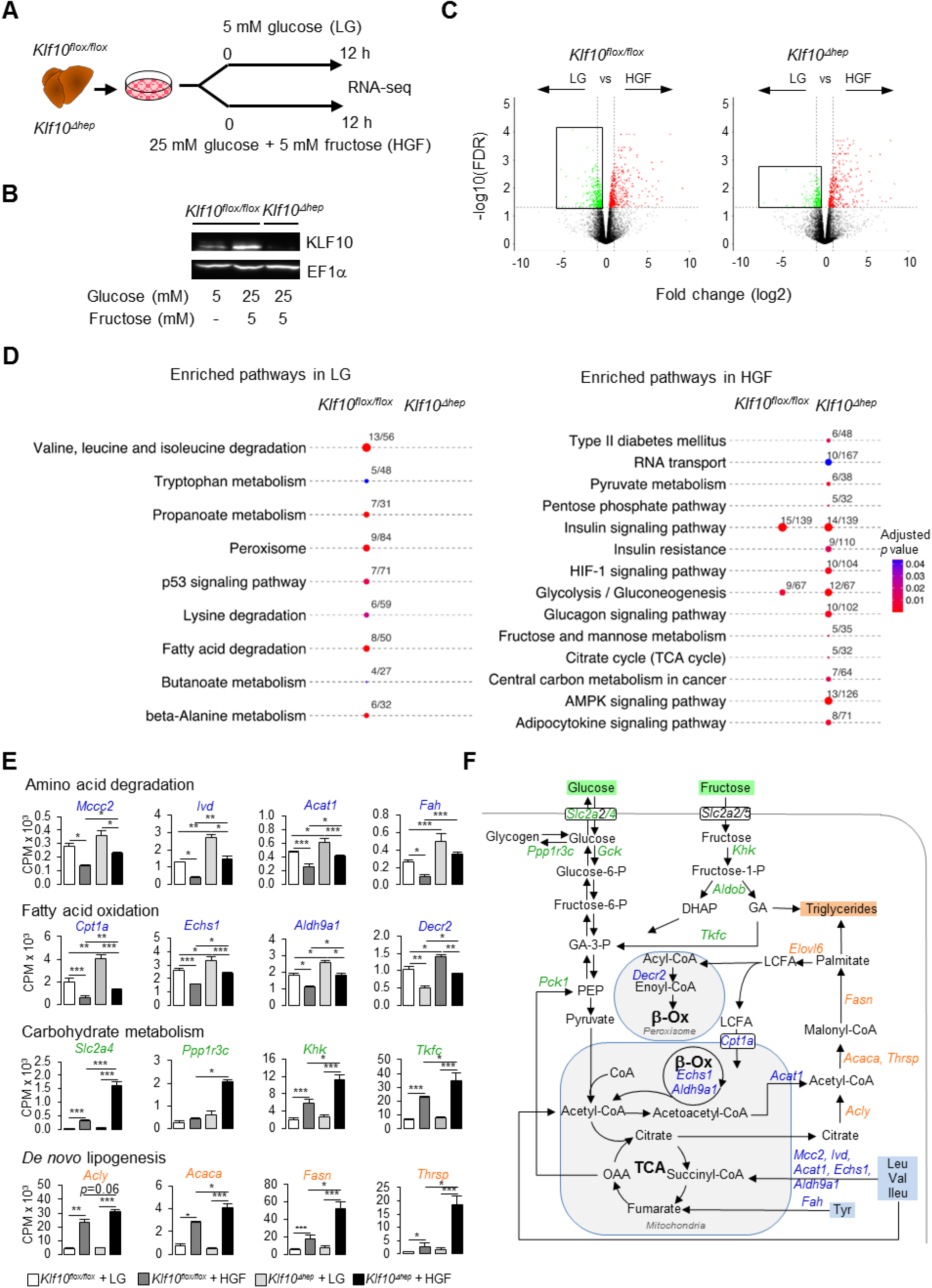
KLF10 governs the transcriptional response to hexose sugars in hepatocytes. (A) Schematic illustrating the design of the *in vitro* sugar challenge in *Klf10*^*flox/flox*^ and *Klf10*^*Δhep*^ hepatocytes treated with low glucose (LG) or high glucose and fructose (HGF). (B) Representative immunoblot showing KLF10 protein abundance in *Klf10*^*flox/flox*^ hepatocytes treated with LG or HGF and in *Klf10*^*Δhep*^ primary hepatocytes treated with HGF. EF1α was used as loading control. (C) Volcano plots showing changes in gene expression in *Klf10*^*flox/flox*^ and *Klf10*^*Δhep*^ hepatocytes treated with LG or HGF. Positive fold change values represent genes upregulated under HGF conditions, negative fold change values represent genes upregulated under LG conditions. Genes significantly upregulated (FDR < 0.5) and displaying a fold change (FC) > 2 in HGF (red dots) and in LG (blue dots) conditions. Dashed horizontal lines, FDR = 0.05; dashed vertical lines, FC = 2. Boxed areas highlight the larger number of downregulated genes in *Klf10*^*flox/flox*^ compared to *Klf10*^*Δhep*^ hepatocytes. (D) KEGG enriched pathways in *Klf10*^*flox/flox*^ and *Klf10*^*Δhep*^ primary hepatocytes treated with LG or HGF. (E) Expression profiles of genes associated with key metabolic pathways in *Klf10*^*flox/flox*^ and *Klf10*^*Δhep*^ primary hepatocytes treated with LG or HGF (mean ± SEM, n = 3). (F) A schematic of key metabolic pathways in hepatocytes. Genes significantly upregulated in *Klf10*^*Δhep*^ vs. *Klf10*^*flox/flox*^ hepatocytes when treated with HGF are highlighted on the schematic. DHAP, dihydoxyacetone phosphate; LCFA, long chain fatty acids; GA, glyceraldehyde; OAA, oxaloacetic acid; PEP, phospho-enol-pyruvate. Statistics: Non parametric Kruskal and Wallis test. *, *p* < 0.05; **, *p* < 0.01; ***, *p* <0.005. See also Figure S5 and Tables S3 and S4.

The switch from the LG to the HGF condition lead to a differential expression of 842 and 608 genes in *Klf10*^*flox/flox*^ and *Klf10*^*Δhep*^ hepatocytes, respectively (Figure 5C, Table S3). Notably, the number of downregulated genes was decreased by ~40% in challenged *Klf10*^*Δhep*^ hepatocytes compared to *Klf10*^*flox/flox*^ hepatocytes, suggesting that the ability of *Klf10*^*Δhep*^ hepatocytes to repress gene expression in response to the HGF challenge is compromised (Figure 5C and S5A). Using the significantly altered transcripts detected under the LG and HGF conditions as an input, we performed pathway enrichment analysis and found several differences between *Klf10*^*flox/flox*^ and *Klf10*^*Δhep*^ hepatocytes (Figure 5C and Table S4). In the LG condition, pathways associated with amino acid catabolism, fatty acid oxidation, and peroxisome metabolism were significantly enriched in *Klf10*^*flox/flox*^ but not *Klf10*^*Δhep*^ hepatocytes (Table S4). Underlying these changes, *Klf10*^*flox/flox*^ hepatocytes displayed an upregulation of genes coding enzymes linked to branched chain amino acids (*Mccc2, Ivd, Acat1, Fah, Echs1, Adlh9a1*), alanine, lysine and tryptophan (*Echs1, Adlh9a1*), and, tyrosine (*Fah*) degradation, as well as genes encoding for enzymes linked to mitochondrial and peroxisomal fatty acid oxidation (*Cpt1a*, *Echs1, Adlh9a1, Decr2*) (Figure 5D, 5E and 5F). In the HGF condition, *Klf10*^*Δhep*^ hepatocytes displayed a greater number of enriched pathways relative to *Klf10*^*flox/flox*^ hepatocytes, with many of these *de novo* pathways associated with carbohydrate metabolism (Figure 5C, right panel). Similar to our results in mice challenged with the chow + SSW diet, the HGF condition resulted in an upregulation of the glucose transporter *Slc2a4* in the *Klf10*^*flox/flox*^ and *Klf10*^*Δhep*^ hepatocytes, with the absence of KLF10 further augmenting this response (Figure 5E). In a similar manner, *Klf10*^*Δhep*^ hepatocytes in the HGF condition displayed greater induction of genes associated with fructose (*Khk, Tfkc*) and glycogen (*Ppp1r3c*) metabolism relative to *Klf10*^*flox/flox*^ hepatocytes (Figure 5D and 5E). In addition to the enhanced upregulation of genes associated with carbohydrate metabolism, *Klf10*^*Δhep*^ hepatocytes had a greater induction of genes associated with *de novo* lipogenesis (*Acly, Acc1, Fasn, Thrsp*) and long chain fatty acid elongation (*Elovl6*) (Figures 5D, 5E, 5F and S5B).

Collectively, our *in vitro* challenge data supports the notion that KLF10 acts as a brake that aids in fine-tuning the hepatocyte’s transcriptional response to glucose and fructose. Furthermore, the dysregulation of various metabolic pathways in hepatocytes lacking KLF10 is consistent with, and may underlie, the abnormal physiological response of *Klf10*^*Δhep*^ mice challenged with the chow + SSW diet.

### KLF10 targets an extensive metabolic gene network in the liver

To further explore the role of KLF10 in the regulation of the hepatic metabolism at the transcriptome level, we assessed the genome-wide occupancy of KLF10 in mouse liver at the time of maximal expression of the protein (ZT9, Figure S6) using chromatin immunoprecipitation followed by high throughput sequencing (ChIP-seq) (Figure 6A). Upon filtering the list of KLF10-bound fragments for the presence of at least one KLF10 binding motif (Schmitges et al., 2016), we obtained a repertoire of KLF10 35,523 KLF10 binding sites present between in the −50 kb and +1 kb region of associated genes (Figure 6B). The density of KLF10 binding sites was negatively correlated with their distance from the transcription initiation site with 37 % being located in the −10 kb and +1 kb, thus indicating that KLF10 mainly binds proximal promoter regions (Figure 6B). We next integrated expression data for genes differentially expressed in *Klf10*^*Δhep*^ vs *Klf10*^*flox/flox*^ hepatocytes in the HGF condition (see Figure 5), ChIP-seq data for genes containing at least one binding site in their −10 kb to +1 kb promoter region and HumanCyc metabolic pathways (Romero et al., 2005) in a single network visualized in Cytoscape. In this network containing 795 nodes and 10,980 edges, we found that genes regulated or/and bound by KLF10 were present in 23 annotated subnetworks (Figure 6C, Table S5). However, half of these genes clustered in three major processes associated with lipid and amino acid metabolism as well as protein phosphorylation (Figure 6C, Cytoscape file S1). The intersect between differentially expressed genes in HGF and KLF10 bound genes identified 93 direct targets (Figure 6D). This list was primarily enriched for genes associated with carboxylic acid metabolism, suggesting a role for KLF10 in central carbon metabolism (Figure 6D). In line with this observation, two of the top ranked direct target genes included *acyl-CoA synthetase short chain family member 2* (*Acss2)* and *acetyl-CoA carboxylase beta* (*Acacb)*, which encode two key enzymes of acetyl-CoA metabolism (Table S5). ACSS2 converts acetate to acetyl-CoA, while ACACB carboxylates acetyl-CoA to malonyl-CoA which is both a negative regulator of fatty acid oxidation through inhibition of CPT1a mediated fatty acid uptake within the mitochondria and the substrate of FASN for palmitate synthesis (Figure 6E). ChIP experiments targeting the KLF10 binding sites identified within the *Acss2* and *Acacb* genes using the ChIP-seq profiling experiment confirmed that both genes were bound by KLF10 at ZT15 in the liver of *Klf10*^*flox/flox*^ mice on chow + SSW diet (Figure 6E). Consistently these two genes showed a greaer upregulation in the liver of *Klf10*^*Δhep*^ mice on chow + SSW diet compared to *Klf10*^*flox/flox*^ mice on the same diet, demonstrating that KLF10 is a transcriptional repressor of these two genes upon induction by high sugar (Figure 6F). Interestingly, ACSS2 is one of the highly connected nodes in the KLF10 metabolic network, suggesting that KLF10 dependent repression of *Acss2* may indirectly impact multiple components of hepatic energy metabolism (Cytoscape file S1 and table S5). We conclude from this data that KLF10 has an extensive repertoire of metabolic targets in the liver that include key regulators of acetyl-CoA metabolism.

**Figure 6.**
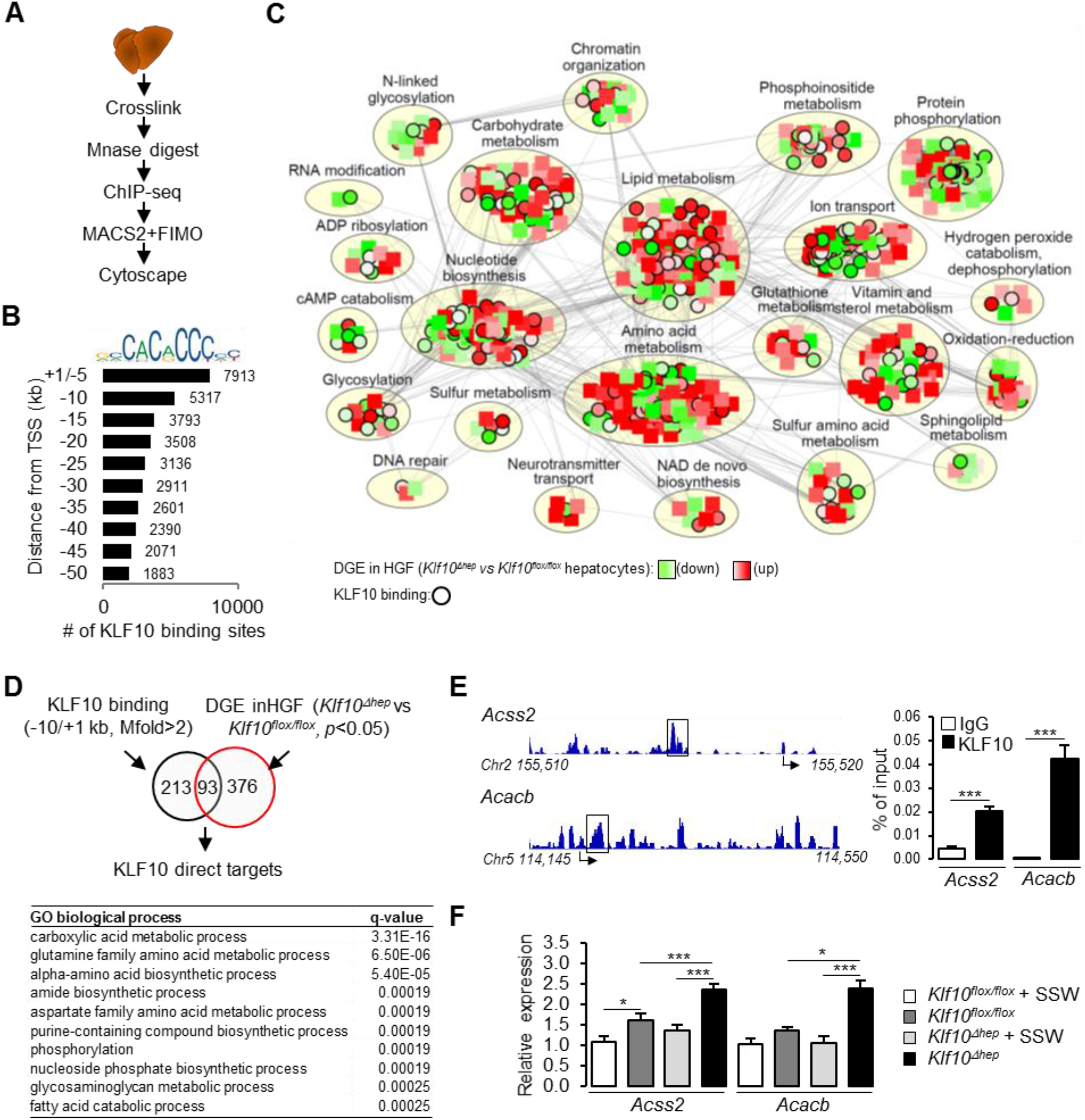
KLF10 regulates a large metabolic network in the liver. (A) Schematic illustrating the workflow used to identify KLF10 bound loci in mouse liver at ZT9. (B) Jaspar logo representing the KLF10 response element DNA sequence and distribution of KLF10 binding sites in mouse liver as a function of distance to the transcription start site. (C) Cytoscape metabolic network integrating RNA-seq data from *Klf10*^*flox/flox*^ and *Klf10*^*Δhep*^ hepatocytes treated with HGF and ChIP-seq data. (D) Venn diagram showing the number of direct KLF10 target (top) and their associated GO biological process annotation (bottom). (E) ChIP-seq tracks near the *Acss2* and *Acacb* promoters (left) and ChIP of the boxed region at ZT15 in the liver of *Klf10*^*flox/flox*^ mice fed the chow + SSW diet (right) (mean ± SEM, n = 4-6). (F) Expression profiles of *Acss2* and *Acacb* at ZT15 in the liver of *Klf10*^*flox/flox*^ and *Klf10*^*Δhep*^ mice given a chow or chow + SSW diet (mean ± SEM, n = 6). Statistics: Non parametric Kruskal and Wallis test. *, *p* < 0.05; **, *p* < 0.01; ***, *p* <0.005. See also Figure S6, Table S5 and Cytoscape file S6.

## Discussion

There is newfound appreciation of KLFs as transcriptional regulators of energy metabolism (Hsieh et al., 2019). This role is underscored by the identification and KLF variants involved in cardiometabolic disorders (Oishi and Manabe, 2018). In the liver, several *Klf* genes are direct targets of the CLOCK transcription factor (Yoshitane et al., 2014), suggesting that timed expression of these transcription factors may help facilitate the connection between the circadian clock and metabolism (Jeyaraj et al., 2012; Panda, 2016; Reinke and Asher, 2019; Sinturel et al., 2020).

In this study, we show that KLF10 is required for the circadian coordination of biological processes associated with energy metabolism in the liver while dispensable for an intact hepatic clock function. This confirms our initial hypothesis that KLF10 acts primarily a clock output (Guillaumond et al., 2010). Alteration of the rhythmic hepatic transcriptome in *Klf10*^*Δhep*^ mice also led to *de novo* oscillation for a large number of genes, reminiscent of the transcriptional changes occurring in KLF15 deficient cardiomyocytes (Zhang et al., 2015, p. 15). In addition to the genetic disruption of clock-controlled KLFs, a variety of other systemic perturbations including high fat and ketogenic diet challenges, arrhythmic feeding, calorie restriction, alcohol consumption, lung cancer and rheumatoid arthritis also generate *de novo* circadian oscillations in the liver (Eckel-Mahan et al., 2013; Gaucher et al., 2019; Greenwell et al., 2019; Makwana et al., 2019; Masri et al., 2016; Poolman et al., 2019; Tognini et al., 2017). Unchallenged *Klf10*^*Δhep*^ mice appear healthy but are prone to develop hepatic steatosis when fed with high sugar while *Klf10^−/−^* mice develop greater liver injury in a non-alcoholic steatohepatitis (NASH) model (Leclère et al., 2020). We therefore interpret the *de novo* oscillation seen in *Klf10*^*Δhep*^ mice as the signature of their hepatic vulnerability. In line with this assumption, we identified apoptosis and inflammation related processes as gene sets gaining circadian coordination in *Klf10*^*Δhep*^ mice. It is presently unknown whether the extensive reprograming of the hepatic circadian transcriptome seen in many genetic or disease mouse models also occurs in humans. Should this occur, it may have important biomedical implications if diagnostic biomarkers or/and regulators of drug pharmacology that are expressed at low levels and arrhythmic in healthy subjects would become circadian upon disease initiation and progression, advocating for the integration of circadian timing in translational research (Cederroth et al., 2019).

Hepatocyte KLF10 furthermore operates on several levels to protect the liver from the negative effects of excess dietary sugars. We found that it suppresses glucose uptake and we obtained correlative evidence that this could be achieved by preventing misexpression of the adipose and muscle specific *Glut4* glucose transporter in hepatocytes. Previous studies showed that *Klf10* is a glucose-responsive gene (Guillaumond et al., 2010; Hirota et al., 2002), a finding which together with our recent observation strongly suggests that KLF10 directs a negative feedback loop limiting excessive glucose intake. Consistent with this role, glycolytic gene expression was upregulated in *Klf10*^*Δhep*^ mice. In addition to glucose, we show that *Klf10* is induced by fructose, and the combination of these two hexose sugars amplifies this response. This upregulation is crucial in blunting the negative effects associated with sugar intake as *Klf10*^*Δhep*^ mice challenged with a sugar-sweetened beverage containing glucose and fructose leads to greater signs of insulin resistance and hepatic steatosis. Our functional genomics data indicates that KLF10 represses the lipogenic gene network in liver. During the metabolic adaptation of the liver to high sugar, the glucose inducible factor carbohydrate-responsive element-binding protein (ChREBP) plays a critical role by inducing lipogenic genes (Ma et al., 2006; Ortega-Prieto and Postic, 2019). Interestingly, the induction of *Klf10* is also mediated by ChREBP (Iizuka et al., 2011). We therefore hypothesize a scenario in which sugar induced steatosis is attenuated by KLF10 *via* a double action taking place upstream and downstream of ChREBP through repression of glucose intake and *de novo* lipogenesis respectively. Further, fructose is also converted by the gut microbiota to acetate which fuels ACLY independent lipogenesis upon conversion to acetyl-coA by liver ACSS2 (Zhao et al., 2020, 2016). Our finding that KLF10 is a transcriptional repressor of *Acss2* suggests an additional mechanism used by KLF10 to reduce the impact of dietary fructose. Our findings are directly relevant to human health as the increased consumption of fructose in the form of sweetened beverages and processed food unarguably contributes to the currently uncontrolled pandemic of obesity and NAFLD (Hannou et al., n.d.; Jensen et al., 2018; Mortera et al., 2019). Links between the circadian clock and NALFD have been suggested yet they are mostly based on preclinical models with disrupted core clock genes, a situation that is unlikely in humans (Mazzoccoli et al., 2018; Mukherji et al., 2019). More presumably, the extensive rewiring of circadian gene expression associated with many pre-disease or disease states and, leaving an intact clock while altering the expression of clock-controlled regulators such as KLF10 may disconnect circadian timing from metabolism and thereby contribute to the development of chronic liver disease including NAFLD.

In summary, our work defines KLF10 as a dual transcriptional regulator with a circadian arm participating in the circadian coordination of liver metabolism and a homeostatic arm serving as a negative feedback control of sugar induced lipogenesis. This duality is likely to be coordinated in healthy hepatocytes so that KLF10 peaks at the fasting/feeding transition when sugar production or intake increases. Desynchrony between circadian timing and feeding as observed in metabolic disorders may therefore challenge KLF10 function with associated negative health outcome.

## Acknowledgements

This study was supported by French Agence Nationale pour la Recherche (ANR): #ANR-15-CE14-0016-01, #ANR-18-CE14-0019-02 and by the “Investments for the Future” LABEX SIGNALIFE (#ANR-11-LABX-0028-01), the UCA^JEDI^ Investments in the Future project (#ANR-15-IDEX-01) and ATER grant from Université Côte d’Azur (JSR). We thank the Canceropôle Provence-Alpes-Côte d’Azur, and the Provence-Alpes-Côte d’Azur Region for the financial support provided to the MetaboCell project. We thank Pierre Chambon for the gift of *SA-CreER*^*T2*^ mice. We thank Aurélie Biancardini, Angelo Barnabeo, Pierre Chérel, Antoine Landouar and the staff of the mouse facility for assistance with the animal care. We thank Sama Rekami for the help with the histopathology and the PFTC facility for technical support.

## Authors contributions

Conceptualization, FD and MT; Project administration, FD; Supervision, FD and MT; Methodology, LM, FD and MT; Investigation, AAR, AGC, SG, LM, JSR, MM, FD and MT; Writing-original draft, AAR, FD and MT; Writing-review and editing, AAR, MS, FD and MT; Resources, MS, SG, LM, FD and MT; Funding acquisition, FD, MS and MT.

## Materials and methods

### Key resources table

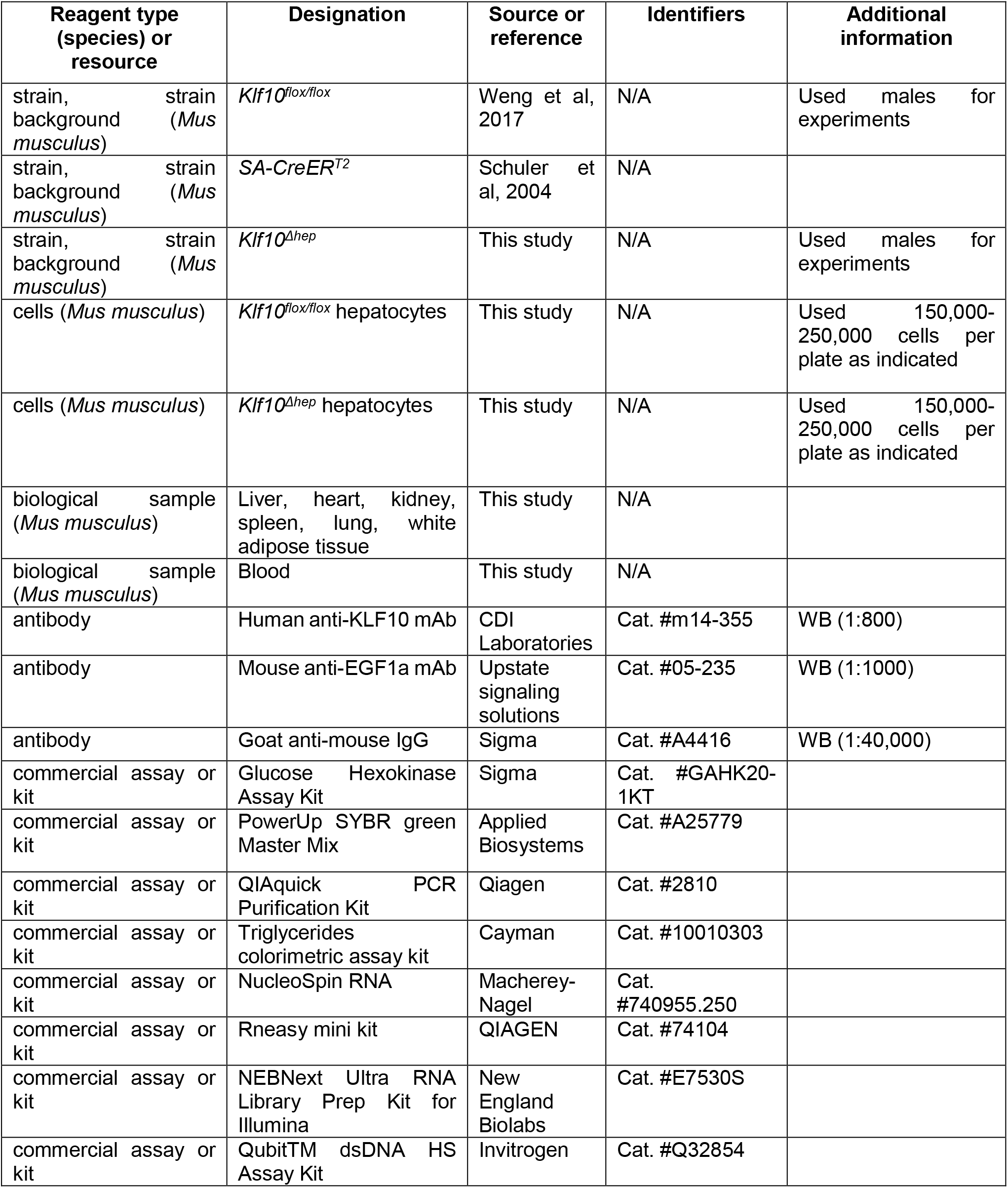
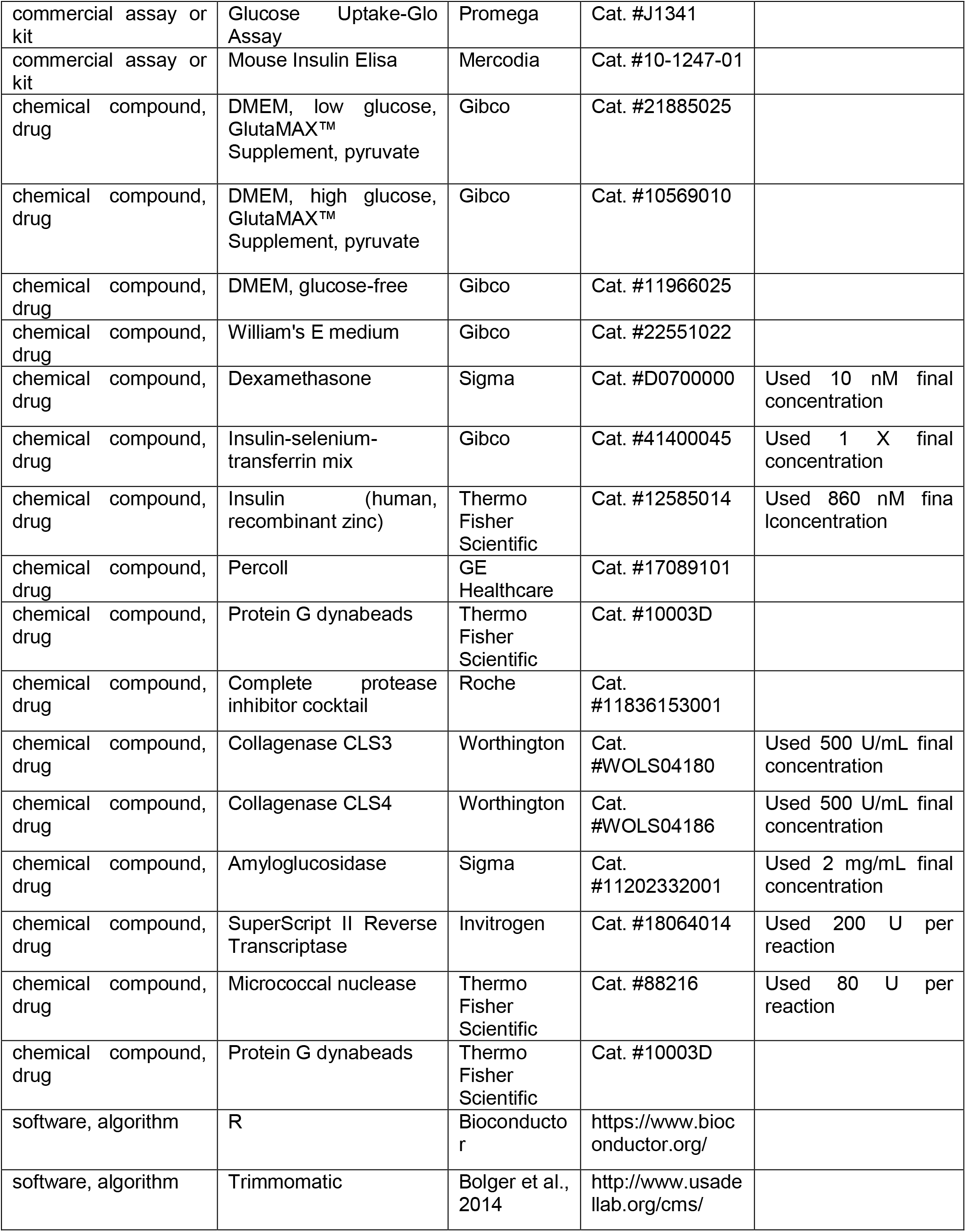
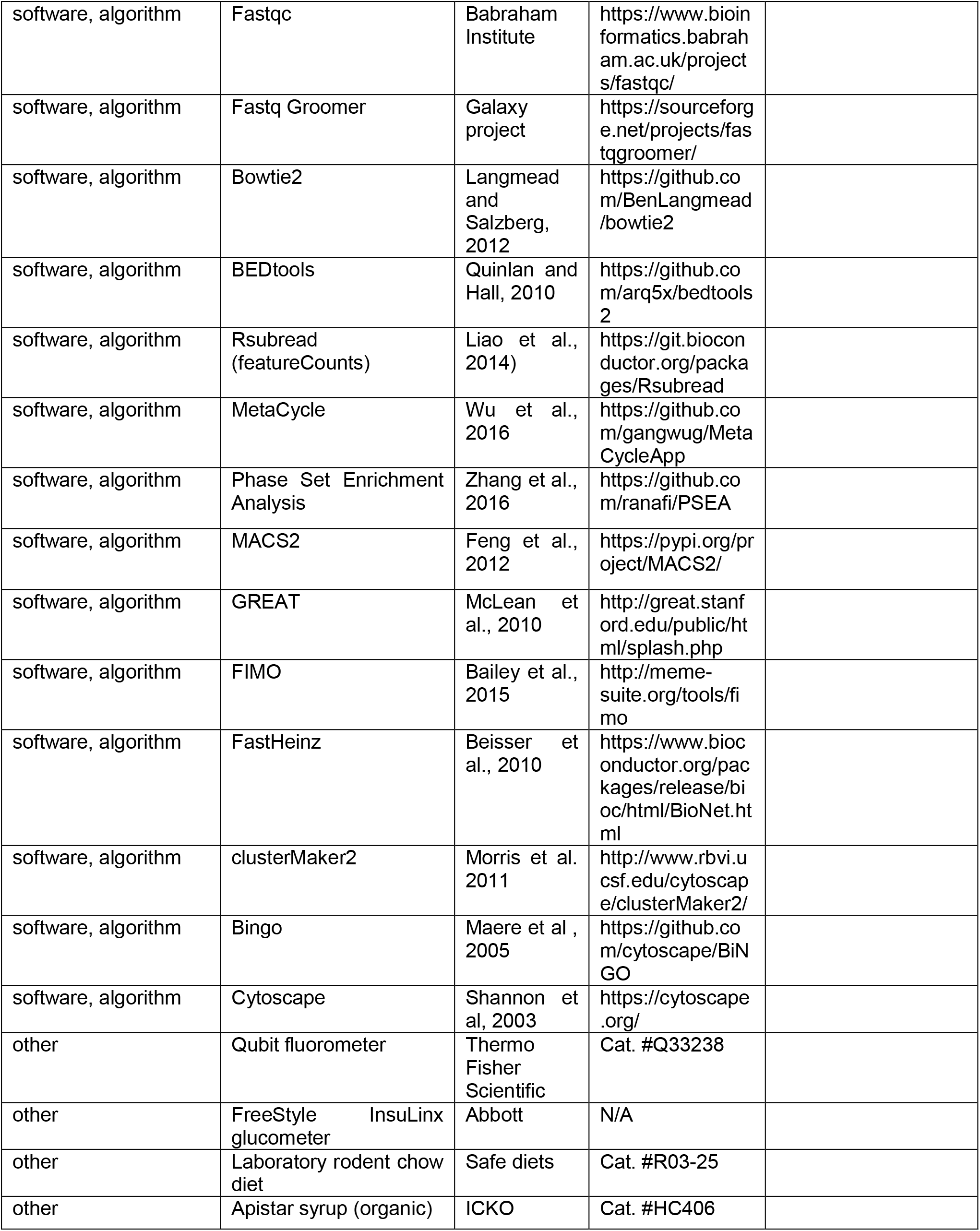

### Mouse models, housing, and diets

To generate a hepatocyte-specific conditional *Klf10* null allele, mice bearing a *Klf10* exon1 flanked by LoxP sites (*Klf10*^*flox/flox*^) described previously (Weng et al., 2017) were crossed with a serum albumin (SA) driven *Cre-ER^T2^* mouse line (Schuler et al., 2004). To obtain mice with a loss of KLF10 in hepatocytes (*Klf10*^*Δhep*^), 5 week old male *Klf10*^*flox/flox*^; *SA*^*+/CreERT2*^ mice were gavaged with tamoxifen (10 mg/kg body weight in 200 μl sunflower oil). For all experiments, aged-matched, tamoxifen treated, *Klf10*^*flox/flox*^ male mice were used as controls. Genotyping was performed by PCR using the primers listed in Supplementary Table S6.

Mice were housed in a temperature- and humidity-controlled room (21±2 °C / 70%), maintained on a 12h:12h LD cycle and fed a standard laboratory chow diet (SafeDiets A03) *ad libitum*. For the *in vivo* sugar challenge, mice from each genotype were randomized between the experimental groups and had *ad libitum* access to standard laboratory chow diet and either water or sugar beverage (water + 30 % Apistar solution [35% fructose, 33% glucose, 32% sucrose]) for 8 weeks. One animal that did not gain weight after eight weeks was excluded.

All animal studies were approved by the local committee for animal ethics Comité Institutionnel d’Éthique Pour l’Animal de Laboratoire (CIEPAL-Azur; Authorized protocols: PEA 244 and 557) and conducted in accordance with the CNRS and INSERM institutional guidelines.

### Primary hepatocyte isolation and culture conditions

Primary hepatocytes were isolated using a retrograde *in situ* perfusion procedure as described previously (Mederacke et al., 2015) with a few modifications. Briefly, mice were anesthetized between ZT3-ZT6, and upon cannulation *via* the inferior *vena cava*, the liver was perfused at a flow rate of 6 mL/min with a warmed (37°C), EGTA solution. After 3 minutes, the buffer was exchanged for a glucose- and EGTA-free buffer containing 45 mM CaCl_2_, collagenase types CLS3 (500 U/mL) and CLS4 (500 U/mL) (Worthington Biochemical Corporation) and the liver was perfused in this buffer for an additional 5 min at a flow rate of 6 mL/min. After perfusion, the liver was excised and placed in Leibovitz L-15 medium supplemented with 10 % fetal calf serum (FCS), 1% penicillin and streptomycin (P/S). Hepatocytes were released by mechanical disruption and filtered through a 70 μm cell strainer and resuspended in 50 mL of Leibovitz L-15, 10% FCS, 1% P/S medium. To seed hepatocytes, the solution was centrifuged at 400 rpm for 2 min and hepatocytes were resuspended in 8 mL William’s E medium supplemented with 860 nM insulin, 1% P/S, 10% FCS, and 1% glutamate. Dead cells were removed by Percoll density-gradient centrifugation. Unless otherwise indicated, hepatocytes were seeded in collagen-coated 6-well plates at a density of 250,000 cells/plate. After hepatocyte attachment, medium was substituted for low glucose DMEM, supplemented with GlutaMAX, pyruvate, 10 nM dexamethasone (Dex), and 1X insulin-selenium-transferrin (ITS). All hepatocyte manipulations took place the following day, ~12-15 h after cell attachment.

### Food seeking behavioural analysis

Mice were individually housed with *ad libitum* access to food and water. An infrared detection system (Actimetrics) was used to assess food seeking behavior. Mice were weighed weekly and their daily food intake was estimated by measuring the difference between the quantity of food provided and food remaining after 1 week. Actogram plots were generated using the ImageJ plugin “ActogramJ” (Schmid et al., 2011).

### Blood analysis

Unless stated otherwise, all measurements were performed at ZT 15. Glycemia levels were measured using a FreeStyle Papillon glucometer (Abbott). Plasma triglycerides were measured using a colorimetric assay kit (Cayman) according to the manufacturer’s protocol. Plasma insulin was measured using a Mouse Insulin Elisa kit (Mercodia) in accordance with the manufacturer’s protocol.

### Glycogen assay

Hepatic glycogen was measured as described previously (Feillet et al., 2016). Hepatic glycogen was extracted from samples collected at ZT3 and ZT15. Upon precipitation, the glycogen pellet was either resuspended in 2 mg/mL amyloglucosidase (Sigma-Aldrich) or in 0.2 M sodium acetate (pH 4.9) to measure total glucose and free glucose, respectively. Total and free glucose was assayed using a glucose-hexokinase assay kit (Sigma-Aldrich) according to the manufacturer’s protocol.

### Glucose tolerance test

Mice were fasted at ZT6 for 6h and then injected intraperitoneally with glucose (0.5g/kg body weight) in 1X PBS. Blood glucose was measured from tail-blood microsamples (<0.5 μL) just before injection and at regular intervals over 2 h using a FreeStyle Papillon glucometer (Abbott).

### Liver triglycerides

Total lipids were extracted from liver samples collected at ZT15 using a modified Folch method (Folch et al., 1957). Briefly, 50-100 mg of liver were homogenized in a methanol/chloroform (2:1, v/v) solution. After addition of 1 volume of chloroform and 0.9 volume of water followed by mixing, the organic phase was separated, evaporated and resuspended in NP40. Triglycerides content was measured using a colorimetric assay kit (Cayman) according to the manufacturer’s protocol.

### Histology

Liver tissue was excised, fixed with 4 % PFA solution overnight, processed for paraffin embedding, sectioned at 5 μm thickness, and then stained for hematoxylin and eosin (H&E). Liver sections were then imaged using the Vectra Polaris digital slide scanner (CLS143455, Akoya Biosciences Inc) and subjected to blind scoring.

### Glucose uptake

Primary hepatocytes from *Klf10*^*flox/flox*^ and *Klf10*^*Δhep*^ were seeded in collagen-coated 24-well plates at a density of 150,000 cells per well and incubated overnight in DMEM containing 25 mM glucose, 17 nM ITS and 10 nM Dex. The following day, cells were washed twice with 1x PBS before stimulation with a glucose-free DMEM containing 17 nM ITS and 1 μM 2-deoxyglucose (2-DG). Hepatocytes were left in this medium for 10 min before the reaction was stopped. Lysates were then assayed for glucose uptake using the Glucose Uptake-Glo Assay kit (Promega) as per the manufacturer’s protocol. Relative light unit (RLU) values were normalized to the total protein concentration of each corresponding sample.

### Liver extract metabolite identification

Liver metabolites were extracted following a protocol adapted from Hui et al., 2017. Briefly, 10 μL/mg liver of −20°C methanol:acetonitrile:water (40:40:20, v/v/v) solution was added to approximately 50 mg of liver. The mixture was crushed using a pellet pestle for 30 s, followed by sonication for 5 min and vortexing for 15 s. Samples were incubated at 4°C for 10 min to precipitate proteins before centrifugation at 4°C and 15,000 *g* for 10 min. The supernatant was transferred to LC-MS vials for analyses.

Liver crude organic extracts were analyzed using a Vanquish UHPLC coupled with a Thermo Q-Exactive (Thermo Fisher Scientific GmbH, Bremen, Germany) mass spectrometer (MS) and an ESI source operated with Xcalibur (version 2.2, ThermoFisher Scientific) software package. Metabolites separation was achieved on a Waters Acquity BEH C8 (1 x 150 mm, 1.7 *μ*m) column with an injection volume of 5 *μL* and a flow rate of 0.1 mLmin^−1^. The mobile phase was composed of octylammonium acetate 4 mM in H_2_O adjusted to pH = 4.6 with acetic acid (phase A) and methanol (phase B) and the following gradient was used: 1% B for 5 min, 1 to 95 % B in 10 min, 95 to 99% B in 5 min, 99% B held for 3 min, then 1% B for 7 min for a total run time of 30 minutes. Data acquisition was realized under full scan switch (positive and negative) mode ionization from *m/z* 80 to 900 at 70,000 resolution followed by inclusion list dependent MS/MS. Samples were randomized and analyzed in triplicates. A quality control (QC) sample was added by pooling equal volumes of each sample to monitor suitability, repeatability and stability of the system and injected every 10 samples. An extraction blank sample was also injected every 10 runs in order to monitor sample cross-contamination. Pure commercial standards (Fumaric acid, succinic acid, oxaloacetic acid, alpha-ketoglutaric acid, glutamic acid, phosphoenolpyruvic acid, dihydroxy acetone phosphate, citric acid, NADHP and pyruvic acid, Merk) were analyzed to monitor their retention time, *m/z* of the parent ion and fragment ions. Feature extraction, compound identification and retention time correction were performed using Compound Discoverer 2.1 software (Thermo Fisher Scientific). Processing of the filtered feature table was realized using Metaboanalyst (www.metaboanalyst.ca) (Chong et al., 2019). Samples were normalized using QCs, data were log transformed then Pareto-scaled (mean-centered and divided by the square root of the standard deviation of each variable). The QC were then removed from the normalized data table to perform statistical analyses.

### *In vitro* stimulation of hepatocytes with glucose and fructose

After seeding of *Klf10*^*flox/flox*^ and *Klf10*^*Δhep*^ hepatocytes, medium was replaced with DMEM supplemented with GlutaMAX, pyruvate, 10 nM Dex, 1X ITS and containing either (i) 5 mM glucose; (ii) 25 mM glucose; (iii) 5mM glucose and 5mM fructose; or (iv) 25 mM glucose and 5 mM fructose. After a 12 h incubation period, media was aspirated and hepatocytes were lysed with Qiagen RLT buffer. Lysed material containing RNA was collected, flash frozen and stored at −80°C until RNA extraction.

### RNA isolation and quantitative real time PCR

Total RNA was purified using RNeasy extraction mini kit (Qiagen) as per the manufacturer’s instructions. To generate cDNAs, 1 μg of total RNA was reverse transcribed using random hexamer primers and Superscript II reverse transcriptase (Invitrogen). Diluted cDNAs (1/10) were added to a PowerUp SYBR green master mix (Applied Biosystems) and amplified using a StepOnePlus Real-Time PCR system (Applied Biosystems) as per the manufacturer’s instructions. The relative mRNA abundance was calculated using the ΔCt method and normalized to the value of the expression of the housekeeping gene *36B4*. The sequences of primers used are listed in supplementary Table S6.

### Western blotting

Whole cell extracts from liver tissue or isolated hepatocytes were processed using the Active Motif Nuclear Extraction kit as per manufacturer’s instructions. Lysates were separated by SDS–PAGE and transferred to PVDF membranes (Bio-Rad). Immunoreactive protein was detected using ECF reagent (Millipore) and Fusion FX7 (Vilber). The primary antibodies used were human anti-KLF10 mAb (1:800), and mouse anti-EGF1a mAb (1:1000). The secondary antibody used was goat anti-mouse IgG (1:40000).

### RNA-sequencing

For the circadian time course experiment, equal amounts of total RNA extracted from 3 mouse livers were pooled at each of the 8 time points assessed over a 24h period. Preparation of libraries and sequencing was performed at UCAGenomiX (Sophia-Antipolis, France). Here, libraries were prepared using the Truseq Stranded Total RNA Preparation Kit with Ribo-Zero Gold (Illumina) following the manufacturer’s recommendations. Normalized libraries were multiplexed, loaded on a flow cell (Illumina) and sequenced on a NextSeq 500 platform (Illumina) using a 2×75bp paired-end (PE) configuration. Image analysis and base calling were conducted by the HiSeq Control Software (HCS). Raw sequencing data (.bcl files) generated from Illumina NextSeq 500 platform was converted into fastq files and de-multiplexed using Illumina’s bcl2fastq 2.17 software.

For the *in vitro* sugar challenge experiment, RNA was extracted from *Klf10*^*flox/flox*^ and *Klf10*^*Δhep*^ hepatocytes treated with either 5 mM glucose or 25 mM glucose and 5 mM fructose for 12 hours (3 biological replicates per experimental condition). Preparation of libraries and sequencing was performed at GENEWIZ (Leipzig, Germany). Here, libraries were prepared using NEBNext Ultra RNA Library Preparation Kit for Illumina (New England Biosystems) following the manufacturer’s recommendations. Normalized libraries were multiplexed, loaded on a flow cell (Illumina) and sequenced on a HiSeq 4000 platform (Illumina) using a 2×150 PE configuration. Image analysis and base calling were conducted by the HiSeq Control Software. Raw sequencing data (.bcl files) generated from Illumina HiSeq 4000 platform was converted into fastq files and de-multiplexed using Illumina’s bcl2fastq 2.17 software.

### Chromatin immunoprecipitation

Snap frozen liver (~300 mg) was cross-linked with 1.1% formaldehyde in PBS for 20 min at room temperature and cross-linking was quenched with 125 mM glycine for 5 min. After Dounce homogenization, the crushed tissue was washed twice in cold PBS, resuspended in cold cell lysis buffer (50 mM HEPES [pH 8], 200 mM NaCl, 20 mM EDTA, 12% glycerol, 1% NP40, 0.8% Triton-X100, 1 mM PMSF, protease inhibitor cocktail), and sonicated 3x (15 sec on/30 sec off cycle, low power mode) in a Bioruptor (Diagenode). Nuclei were purified by sedimentation at 1,000 rpm for 5 min, the supernatant was removed and the pellet was resuspended in nuclei preparation buffer (0.34 M sucrose, 15 mM Tris [pH 8], 15 mM KCl, 0.2 mM EDTA, 0.2mM EGTA, protease inhibitor cocktail). To digest chromatin, the suspension was incubated with 80 Units micrococcal nuclease (MNase) for 10 min at 37°C. Next, the suspension was spun at 6,600 rpm for 5 min, the supernatant was discarded, and nuclei were resuspended in nuclei lysis buffer (50 mM Tris [pH8], 150 mM NaCl, 10 mM EDTA, 0.5% NP40, 1% Triton-X100, 0.4% SDS and protease inhibitor cocktail). The sample was sonicated for 12x (30 sec on/30 sec off, high power mode) in a bioruptor to release digested chromatin. The suspension containing the fragmented chromatin was centrifuged at 10,000 rpm for 8 min at 4°C to remove debris and the supernatant was diluted in ChIP dilution buffer (50 mM Tris pH8, 150 mM NaCl, 1% Triton-X100, protease inhibitor cocktail). Protein-DNA complexes were immunoprecipitated using 3-4 μg of human anti-KLF10 mAb (CDI laboratories) incubated at 4°C overnight and subsequently isolated with protein G Dynabeads for 2 h at room temperature. Beads were washed 4 times in low salt buffer (50 mM Tris [pH7.4], 5 mM EDTA, 150 mM NaCl, 1% Triton X-100, 0.5% NP40), 4 times in high salt wash buffer (100 mM Tris [pH7.4], 250 mM NaCl, 1% NP40, 1% sodium deoxycholate), once in TE buffer (50 mM Tris [pH7.4], 10 mM EDTA) and then eluted with 1% SDS-0.1 M NaHCO_3_. Reverse crosslinking of protein-DNA complexes was performed overnight by incubation at 65 °C followed by proteinase K digestion for 4 h at 42 °C. DNA fragments were purified using QIAquick PCR purification columns (Qiagen) quantified using a Qubit fluorometer (Invitrogen). Fragmented DNA was flash frozen and stored at −80°C until further processing for high throughput sequencing or qRT-PCR.

### ChIP-sequencing

Preparation of libraries and sequencing was performed at GENEWIZ (Leipzig, Germany). Here, libraries were prepared using NEBNext Ultra DNA Library Preparation Kit for Illumina (New England Biosystems) following the manufacturer’s recommendations. Briefly, the ChIP DNA was end repaired and adapters were ligated after adenylation of the 3’ends. Adapter-ligated DNA was size selected, followed by clean up, PCR enrichment. ChIP libraries were validated using an Agilent TapeStation, and quantified using Qubit 2.0 Fluorometer as well as qRT-PCR (Applied Biosystems, Carlsbad, CA, USA). Normalized libraries were multiplexed, clustered, and loaded on one lane of a flow cell (Illumina). Libraries were sequenced on a HiSeq 4000 platform (Illumina) using a 2×150 PE configuration. Image analysis and base calling were conducted by the HiSeq Control Software (HCS). Raw sequence data (.bcl files) generated from Illumina HiSeq was converted into fastq files and de-multiplexed using Illumina’s bcl2fastq 2.17 software. One mis-match was allowed for index sequence identification.

### Network analysis

A Network integrating RNA-seq data from the HGF condition of the *in vitro* sugar challenge experiment and ChIP-seq data was generated using FastHeinz from the R BioNet package (Beisser et al., 2010) with the HumanCyc metabolic pathways (http://humancyc.org/) downloaded from http://www.ndexbio.org (Pillich et al., 2017). Next, a weight matrix was constructed with all expressed genes in the *Klf10*^*flox/flox*^ and *Klf10*^*Δhep*^ primary hepatocytes taking the lowest *p* value from the ChIP-seq RNA-seq datasets. Network visualization was done using Cytoscape and AutoAnnotate app (Kucera et al., 2016; Shannon et al., 2003). Clustering was performed using the MCL partitioning algorithm from clusterMaker2 (Morris et al., 2011) and gene ontology analysis was performed using BiNGO (Maere et al., 2005) at http://impala.molgen.mpg.de/ (Kamburov et al., 2011) and at https://www.uniprot.org/.

### Quantification and statistical analyses

For the circadian time course experiment, sequences were preprocessed using FASTQ groomer and checked using Fastqc (https://www.bioinformatics.babraham.ac.uk/projects/fastqc/). Reads were mapped against the *Mus musculus* genome (mm10) downloaded from Ensembl.org using STAR_2.4.0a (Dobin et al., 2013) with the Encode RNA-seq best practices options indicated (“– outFilterIntronMotifsRemoveNoncanonicalUnannotated –alignMatesGapMax 1000000 – outReadsUnmapped Fastx –alignIntronMin 20 –alignIntronMax 1000000 –alignSJoverhangMin 8 –alignSJDBoverhangMin 1 –outFilterMultimapNmax 20”). After alignment to the genome reads were counted with featureCounts (Liao et al., 2014) with “–primary –g gene_name –p –s 1 –M –C” options indicated. Gene stable IDs (GRCm38.p6) were downloaded from Ensembl Genes 94 (Ensembl) and used to filter for mouse protein coding genes. Matrix of raw counts from both genotypes across time was generated in R (R Core Team, 2018). Transcripts with less than 8 counts across all samples were excluded from downstream analyses. All transcript counts were normalized using edgeR (Robinson et al., 2010) and expressed in counts per million (cpm).

For the *in vitro* sugar challenge experiment, sequences were preprocessed using FASTQ groomer and checked using Fastqc (https://www.bioinformatics.babraham.ac.uk/projects/fastqc/). Reads were mapped against the *Mus musculus* genome (mm10) downloaded from Ensembl.org using HISAT2with default parameters (Kim et al., 2015). After alignment to the genome reads were counted with featureCounts (Liao et al., 2014) with with default parameters. Matrices containing the counts were loaded into R. For downstream analyses, we analyzed protein coding genes with a sum >20 counts across all samples.

Circadian expression of mRNA was determined using MetaCycle (Wu et al., 2016) with the following options: minper = 24, maxper = 24, cycMethod = c (“ARS,” “JTK,” “LS”), analysisStrategy = “auto,” outputFile = TRUE, outIntegration = “both,” adjustPhase = “predictedPer,” combinePvalue = “fisher,” weightedPerPha = TRUE, ARSmle = “auto,” and ARSdefaultPer = 24. Transcripts with an integrated p-value (meta2d_pvalue) < 0.05 were considered rhythmically expressed.

Circadian pathways were determined by Phase Set Enrichment Analysis (PSEA) (Zhang et al., 2016) based on the sets of circadian transcripts with a relative amplitude value (rAMP; the ratio between amplitude and baseline expression) of ≥ 0.1. Gene sets were downloaded from the Molecular Signatures database (MSigDB) H (HallmarkGene Sets) (Subramanian et al., 2005). Sets containing fewer than 5 circadian transcripts were excluded from the analysis. The Kuiper test was used to identify circadian gene sets by comparing the acrophases of all circadian transcripts belonging to each gene set to a uniform background distribution and by testing for non-uniformity (q < 0.01).

For the *in vitro* sugar challenge experiment, differential expression analysis between conditions was performed using the TestDE function in the R package “edgeR” (Robinson et al., 2010). Genes with an FDR value < 0.05 when comparing LG vs HGF conditions in each respective genotype were considered differentially expressed and used to assess the pathways enriched under the LG and HGF conditions. Gene lists were uploaded to enrichR web server (Chen et al., 2013; Kuleshov et al., 2016) and pathway enrichment was assessed by comparing the overlap of genes in the query to the Kyoto Encyclopedia of Genes and Genomes (KEGG) 2019 mouse gene sets (Kanehisa and Goto, 2000). For a schematic of the workflow used, refer to Figure S5C.

For ChIP-seq read mapping, peak calling and quantification, sequences were preprocessed using FASTQ groomer and checked using Fastqc (https://www.bioinformatics.babraham.ac.uk/projects/fastqc/). PE reads were trimmed to 50 bases with Trimmomatic (Bolger et al., 2014), and reads were aligned to mouse genome (GRCm38/mm10) using Bowtie2 (Langmead and Salzberg, 2012) with the default parameters. Next, peak calls were made in MACS2 (Feng et al., 2012) with default parameters, except allowing for duplicates. Filters were applied to keep peaks with a log *p*-value < −2 and a Mfold > 2. Results from the independent ChIP-seq runs were concatenated and intervals were merged using the MergeBED function from BEDtools (Quinlan and Hall, 2010). Identified peaks within the −10 kb to +1 kb region relative to the TSS were annotated using GREAT (McLean et al., 2010) and further filtered for the presence of a KLF10 motif (matrix ID: MA1511.1, http://jaspar.genereg.net/) using FIMO (Bailey et al., 2015).

Numerical values are presented as mean ± SEM. Replicates (n) are indicated in the figure legends. Pairwise comparisons were tested using the non-parametric Wilcoxon test. Differences between more than 2 experimental conditions were tested using the Kruskal and Wallis rank sum test for multiple comparison followed by a pairwise *post-ho*c test using the Benjamini-Hochberg adjustment. *P* values less than 0.05 were considered significant.

## Resource and data availability

Further information and requests for resources and reagents should be directed to and will be fulfilled by the Lead Contact, Michèle Teboul.

The datasets generated during this study are available at the European Nucleotide Archive (ENA) (https://www.ebi.ac.uk/ena/browser/home) under ENA project numbers PRJEB39035, PRJEB39036, PRJEB40195.

## Supplemental files

**Table S1.** Hepatic circadian transcriptome of *Klf10*^*flox/flox*^ and *Klf10*^*Δhep*^ mice

Related to Figure 1. Metacycle analysis of the hepatic transcriptome of *Klf10*^*flox/flox*^ and *Klf10*^*Δhep*^ mice fed *ad libitum* and entrained in a 12:12 LD cycle. (.xlsx 9.65 MB)

**Table S2.** Temporal coordination of the hepatic transcriptome of *Klf10*^*flox/flox*^ and *Klf10*^*Δhep*^ mice

Related to Figure 1. PSEA analysis of the hepatic transcriptome of *Klf10*^*flox/flox*^ and

*Klf10*^*Δhep*^ mice fed *ad libitum* and entrained in a 12:12 LD cycle. (.xlsx 0.02 MB)

**Table S3**. Transcriptome of *Klf10*^*flox/flox*^ and *Klf10*^*Δhep*^ hepatocytes challenged with high sugar

Related to Figure 5. Lists of differentially expressed genes in *Klf10*^*flox/flox*^ and *Klf10*^*Δhep*^ hepatocytes challenged with high sugar compared to unchallenged cells. (.xlsx 0.13 MB)

**Table S4**. Enriched pathways in *Klf10*^*flox/flox*^ and *Klf10*^*Δhep*^ hepatocytes challenged with high sugar

Related to Figure 5. Lists of KEGG enriched pathway in *Klf10*^*flox/flox*^ and *Klf10*^*Δhep*^ hepatocytes challenged with high sugar compared to unchallenged cells. (.xlsx 0.08 MB)

**Table S5.** KLF10 regulated network in mouse liver.

Related to Figure 6. List of genes bound by KLF10 in their −10/+1kb region or differentially expressed in *Klf10^flox/flox^ vs Klf10^Δhep^* hepatocytes challenged with high sugar. (.xlsx 0.16 MB)

**Table S6**. List of Primers used in the study (.xlsx 0.07 MB)

**Cytoscape file S1.** KLF10 regulated metabolic network

Ralated to Figure 6. File used to generate, visualize and analyze the KLF10 regulated metabolic network using Cytoscape (.cys 0.47 MB)

## Supplemental Figures

**Figure S1 (related to Fig. 1).**
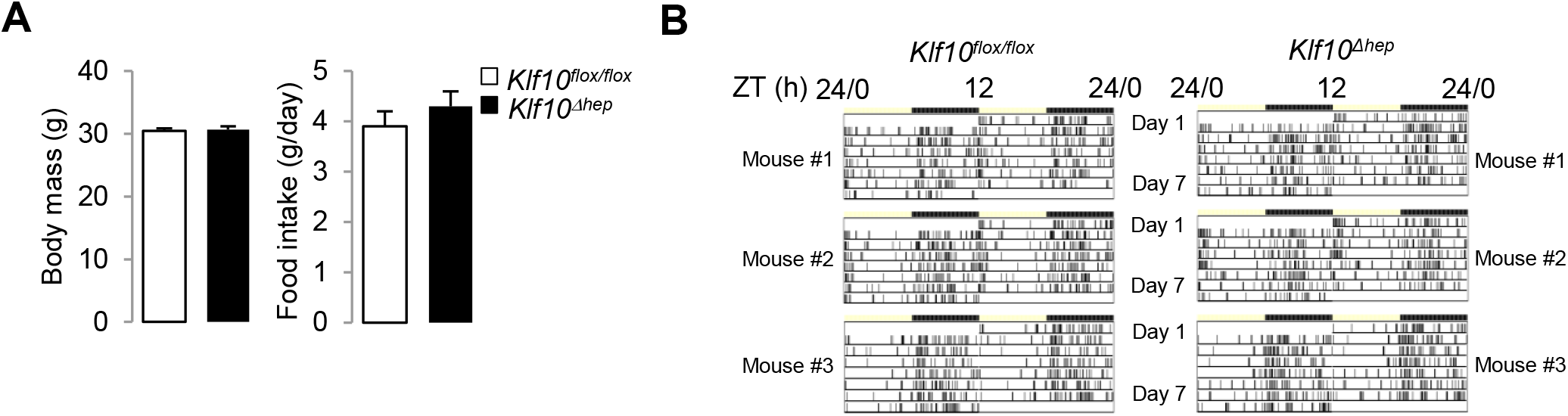
Genetic disruption of *Klf10* in mouse hepatocytes. (A) Body weights of *Klf10*^*flox/flox*^ and *Klf10*^*Δhep*^ mice (mean ± SEM, n = 6). (B) Daily food intake of of *Klf10*^*flox/flox*^ and *Klf10*^*Δhep*^ mice (mean ± SEM, n = 6). (C) Representative actograms of double-plotted food-seeking activity records of *Klf10*^*flox/flox*^ and *Klf10*^*Δhep*^ mice. Mice were placed in individual cages containing a Minimetter infrared detection system. Vertical bars represent time points that animals were in the area of the cage containing food.

**Figure S2 (related to Fig. 2).**
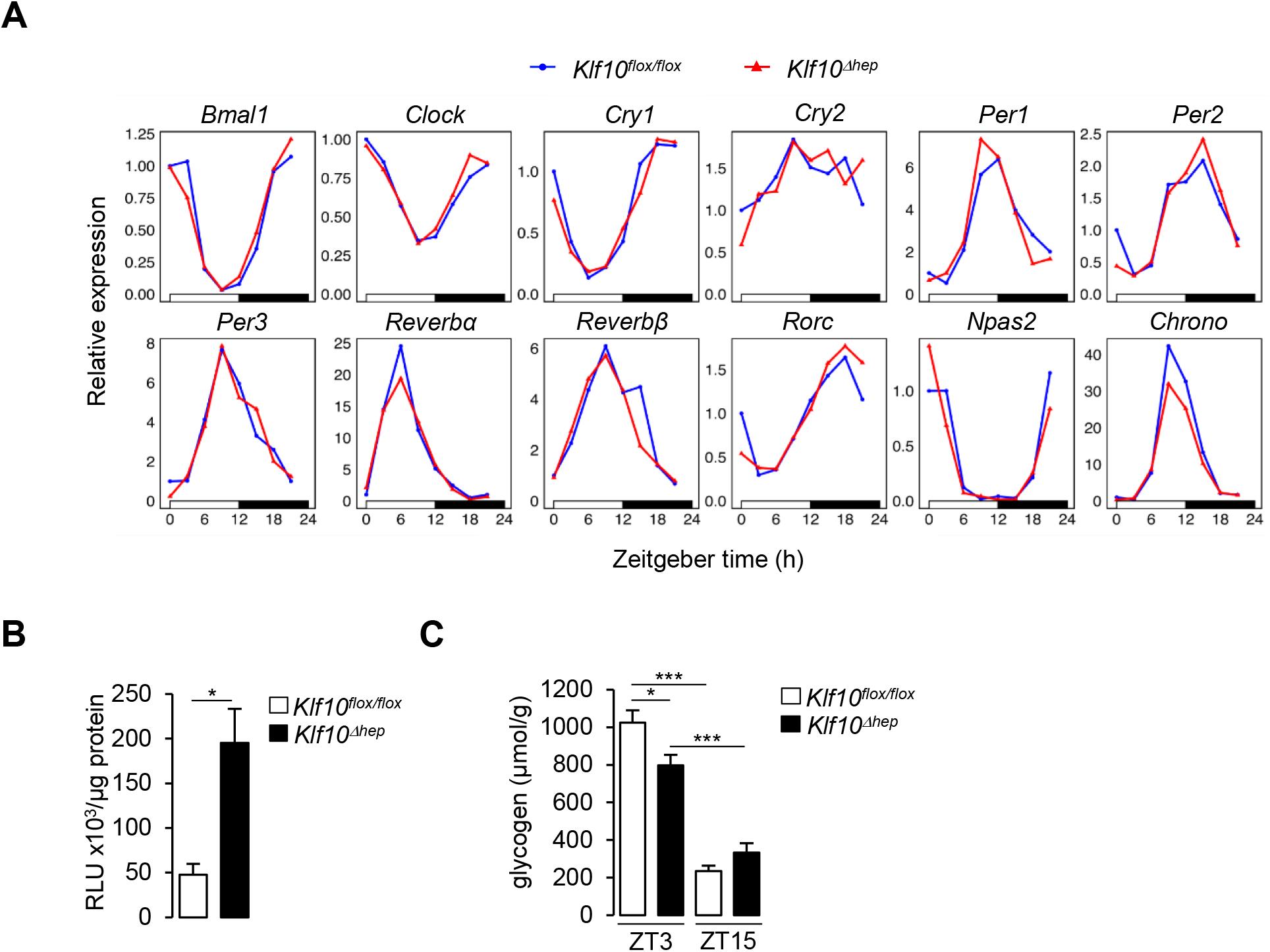
Deletion of hepatocyte KLF10 alters the circadian transcriptome in the liver. (A) Gene expression profiles of clock genes in the livers of *Klf10*^*flox/flox*^ and *Klf10*^*Δhep*^ mice (n = 3 pools of liver per time point). (B) Glucose uptake in *Klf10*^*flox/flox*^ and *Klf10*^*Δhep*^ mice (mean ± SEM, n = 6). (C) Liver glycogen content in *Klf10*^*flox/flox*^ and *Klf10*^*Δhep*^ mice measured at ZT3 and ZT15 (mean ± SEM, n = 5-6). Statistics: Non parametric Wilcoxon (B) or Kruskal and Wallis © test. * *p* < 0.05; ***, *p* <0.005.

**Figure S3 (related to Fig. 3).**
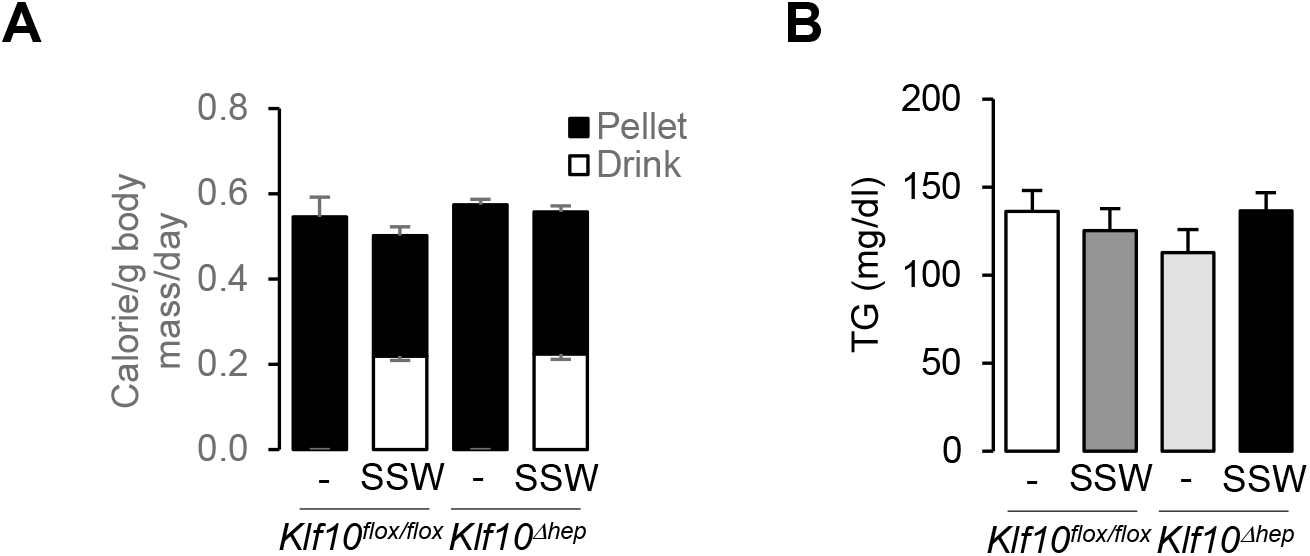
Loss of hepatocyte KLF10 exacerbates the adverse effects associated with increased sugar consumption. (A) Daily caloric intake in *Klf10*^*flox/flox*^ and *Klf10*^*Δhep*^ mice under the chow and chow + SSW dietary conditions (mean ± SEM, n = 8-12). (B) Blood triglyceride content at ZT15 in *Klf10*^*flox/flox*^ and *Klf10*^*Δhep*^ mice under the chow and chow + SSW dietary conditions (mean ± SEM, n = 7-9).

**Figure S4 (related to Fig.4).**
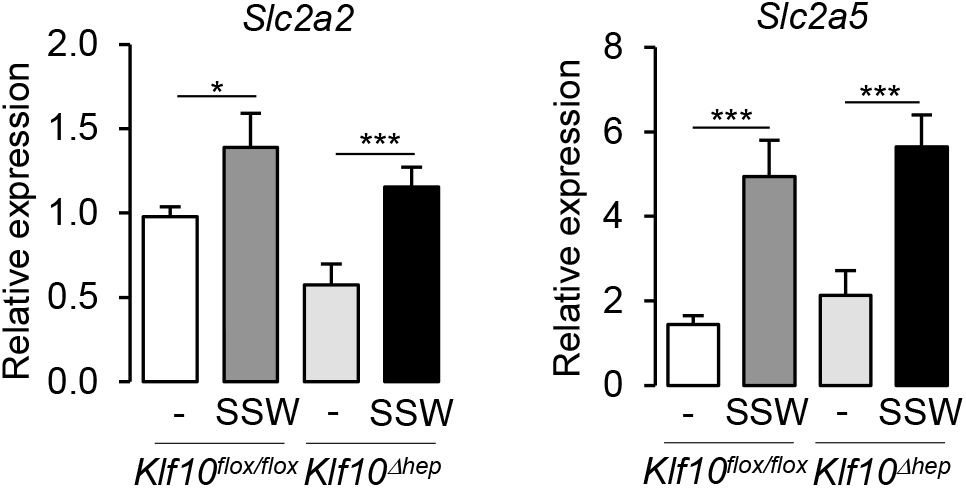
Altered metabolic gene expression in *Klf10*^*Δhep*^ mice challenged with high sugar. Expression profiles of the genes encoding for facilitated glucose transporter *Slc2a2* and the facilitated glucose/fructose transporter *Slc2a5* at ZT15 in *Klf10*^*flox/flox*^ and *Klf10*^*Δhep*^ mice given a chow or chow + SSW diet (mean ± SEM, n = 6). Statistics: Non parametric Kruskal and Wallis test. *, *p* < 0.05; ***, *p* <0.005.

**Figure S5 (related to Fig. 5).**
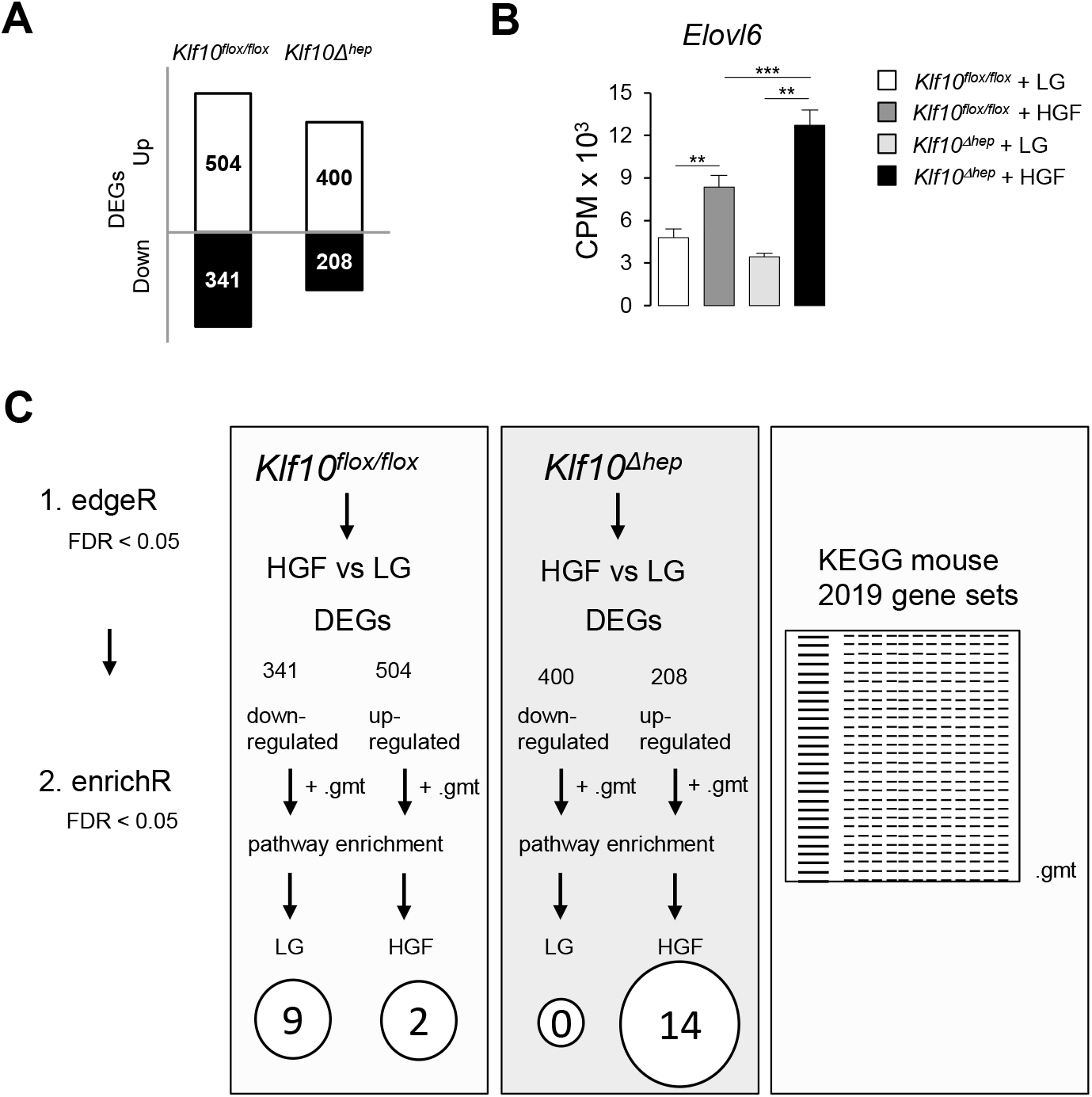
KLF10 governs the transcriptional response to hexose sugars in hepatocytes. (A) Total number of DEGs in *Klf10*^*flox/flox*^ and *Klf10*^*Δhep*^ hepatocytes treated with LG or HGF for 12 hours. (B) Expression profiles of *Elovl6* in *Klf10*^*flox/flox*^ and *Klf10*^*Δhep*^ primary hepatocytes treated with LG or HGF (mean ± SEM, n = 3). (C) Schematic of the workflow used for the assessment of enriched gene sets in *Klf10*^*flox/flox*^ and *Klf10*^*Δhep*^ hepatocytes treated with LG or HGF. A data table containing the enrichment results (input set of genes overlapping with ‘KEGG mouse 2019 gene sets’) was downloaded from the EnrichR server. Gene sets with a q value (FDR) of < 0.05 were used for the dot plot visualization of enriched pathways shown in Figure 3C. Statistics: Non parametric Kruskal and Wallis test. **, *p* < 0.01; ***, *p* <0.005.

**Figure S6 (related to Fig. 6).**
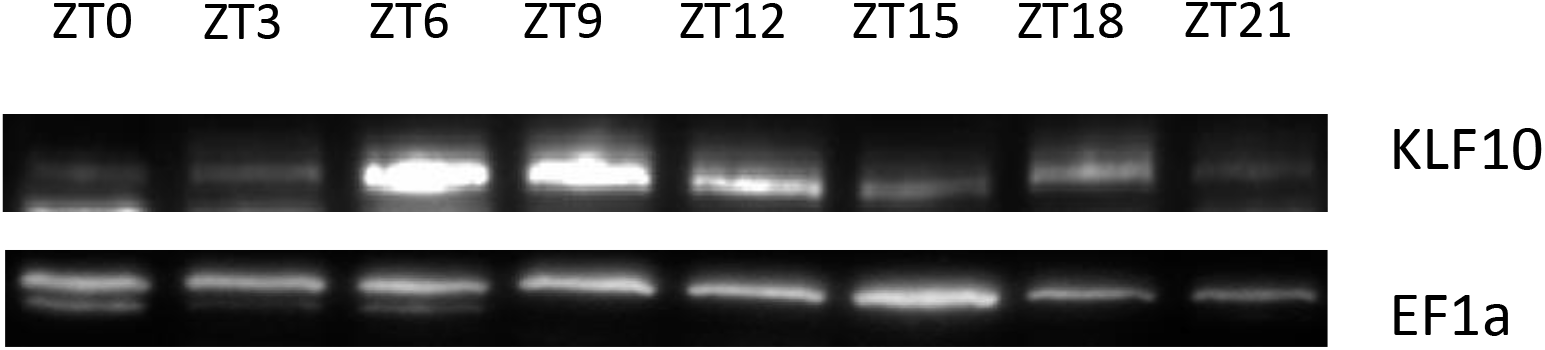
KLF10 regulates a large metabolic network in liver. Immunoblot showing the oscillation of KLF10 protein abundance in the liver of mice entrained in a LD12:12 light/dark cycle.

